# ER-to-Golgi trafficking *via* a dynamic intermediate *cis-*Golgi tubular network in Arabidopsis

**DOI:** 10.1101/2023.10.27.563925

**Authors:** Louise Fougère, Magali Grison, Patricia Laquel, Matheus Montrazi, Fabrice Cordelières, Mónica Fernández-Monreal, Christel Poujol, Tomohiro Uemura, Akihiko Nakano, Yoko Ito, Yohann Boutté

**Affiliations:** Laboratoire de Biogenèse Membranaire, Univ. Bordeaux, UMR5200 CNRS, Villenave d’Ornon, France; Université de Bordeaux, CNRS, INSERM, Bordeaux Imaging Center (BIC), US4, UAR 3420, F-33000 Bordeaux, France; Faculty of Core Research, Natural Science Division, Ochanomizu University, Tokyo, Japan; Live Cell Super-Resolution Imaging Research Team, RIKEN Center for Advanced Photonics, Wako, Saitama, Japan; Institute for Human Life Science, Ochanomizu University, Tokyo, Japan

## Abstract

Endoplasmic Reticulum (ER)-to-Golgi trafficking is a central process of the secretory system of eukaryotic cells that ensures proper spatiotemporal sorting of proteins and lipids^1–5^. However, the nature of the ER-Golgi Intermediate Compartments (ERGIC) and the molecular mechanisms mediating the transition between the ERGIC and the Golgi, as well as the universality of these processes amongst Eukaryotes, remain undiscovered. Here, we took advantage of the plant cell system in which the Golgi is highly dynamic and in close vicinity to the ER^6–9^. We discovered that the ERGIC is composed from at least two distinct subpopulations of *cis*-Golgi. A subpopulation is a reticulated tubulo-vesicular network mostly independent from the Golgi, highly dynamic at the ER-Golgi interface and crossed by ER-induced release of luminal cargos at early stage. Another subpopulation is more stable, cisterna-like and mostly associated to the Golgi. Our results identified that the generation and dynamics of the ER-Golgi intermediate tubulo-vesicular network is regulated by the acyl-chain length of sphingolipids as well as the contacts it establishes with existing Golgi cisternae. Our study is a major twist in the understanding of the Golgi by identifying that the ERGIC in plants is a Golgi-independent highly dynamic tubular network from which arise more stable cisternae-like Golgi structures. This novel model presents a mechanism for early secretory trafficking adapted to respond to developmental and environmental stimuli, including susceptibility or resistance to diseases, autophagy or cell-reprograming.

## Main

The secretory system is an intricate set of membranes that plays fundamental functions to coordinate fluxes and ensure the accurate destination of molecules involved in cellular processes and responses to developmental and environmental stimuli. The first point in the carriageway of this system occurs at the endoplasmic reticulum (ER)-export sites (ERES) that convey proteins to the Golgi apparatus^9,10^. In animal cells, the Golgi apparatus is a ribbon of the Golgi units gathered at the centrosome-nucleated microtubules. In plant cells, Golgi units are dispersed within the cytosol and move along actin cables close to the ER. In animal cells, the traditional vesicle carriers model was challenged by the recent nano-resolved structure of ERES that was identified as a continuous network of interwoven membrane tubules that connects the ER and emits pearled membrane extensions that lie in the direction of the Golgi apparatus^4^. The ERES-derived transport carriers were previously known to be enriched at the tubulo-vesicular ER-Golgi Intermediate Compartment (ERGIC), but how the ERGIC-to-Golgi transition is ensured remain unadressed^4,11^. In plant cells, due to the close proximity of the Golgi and ERES, it has been suggested that no equivalent of the ERGIC exists^6,12^. However, it has been described that some part of the ER, or some ER-derived protrusions, can from contacts with the Golgi apparatus^7,13–15^. In addition, it has been observed that inducing Golgi movement by trapping the Golgi with optical tweezers results in the dynamic remodeling of the ER network, thereby suggesting that the ER and the Golgi are in physical contact^16^. However, the resolution limit of the microscopes employed in plant cells studies has not been sufficient to conclude that the ER-protrusions are tubular intermediates between the ER and the Golgi. In plant cells, it has been shown that while the medial and *trans* cisternae of the Golgi apparatus disappear upon treatment with the Golgi-denaturing drug brefeldin A (BFA), and their proteins absorbed into the ER, the *cis*-Golgi Qa-SNARE SYP31 and Retrieval protein1B (RER1B) remain located in dotty structures that are in close proximity to the ERES. Those structures, or the core *cis*-cisternae where SYP31 and RER1B localize are called “Golgi Entry Core Compartment (GECCO)”, given their special nature at the Golgi entry face^17^. In plants and yeasts, GECCO is thought to be the equivalent of the animal ERGIC^5,18–20^. However, the identity and morphology of these structures as well as their dynamics and the molecular mechanisms involved in their transition to the Golgi remain unknown. In this study we employed improved resolution airyscan microscopy combined with novel quantification methods to track, detect and quantify potential dynamic association between pre-Golgi and Golgi compartments in native BFA-untreated live cells. Unexpectedly, our results revealed that a subpopulation of *cis*-Golgi labeled by the Qb-SNARE MEMBRIN12 (MEMB12) is made of tubules and vesicles, is mostly independent from the rest of the Golgi and displays a highly dynamic behavior characterized by transient associations with the medial-Golgi as well as with the ERES. We established the Retention Using Selective Hooks (RUSH) system in plants to release cargos synchronously from the ER and showed that luminal cargos are crossing the MEMB12 subpopulation upon release. Importantly, we establish lifetime tau-STimulated Emission Depletion (τ-STED) and Expansion Microscopy (ExM) super-resolution methodologies to decipher the structure of pre-Golgi compartments in plants. Our results identified a Golgi-independent MEMB12-positive tubular network that resembles the Vesicular-Tubular Cluster (VTC)/ERGIC of animal cells and that link locally with the Golgi cisternae. Furthermore, we observed that the *cis*-Golgi marker SYP31 labels a more stable Golgi-associated cisterna. We propose that the highly dynamic Golgi-independent MEMB12-tubular structure is part of the ERGIC in plants and that it constitutes an early structure within the GECCO. Our study further identified a functional regulation of the dynamics of the ER-Golgi intermediate structures by both sphingolipids and local contacts with Golgi cisternae that stabilize the intermediate-network into a larger cisterna-like structure. This study proposes a complete reevaluation of the ER-Golgi interface in plant cells by identifying an ER-to-Golgi trafficking *via* a highly dynamic *cis*-Golgi tubular network and by providing novel structural and functional identification of the ERGIC/GECCO in plants.

### A subpopulation of the *cis*-Golgi is Golgi-independent and highly dynamic

GECCO was first identified in BY-2 tobacco cell culture, upon inhibition of BFA-sensitive ARF-GEFs by BFA treatment^17^. In Arabidopsis, given that the ARF-GEF GNOM-LIKE1 (GNL1) is involved in ER-Golgi transport and resistant to BFA, we introduced the *cis*-Golgi GFP-SYP31 fluorescent marker in *gnl1* knockout mutant in which an engineered GNL1-BFA sensitive version was introduced beforehand^21,22^. In this genetic background, our results indicate that GFP-SYP31 remains in small dotty structures after BFA treatment (**Extended data 1a, b**). Contrastingly, the *trans*-Golgi marker ST-mRFP is relocated to the ER upon BFA (**Extended data 1c, d**). These results indicate that GECCO-like structures exist in the Arabidopsis root model. We next checked whether the *cis*-Golgi could behave, for a portion of it, independently from the rest of the Golgi in native condition, i.e without BFA in wild-type genetic background. To test this hypothesis, we used airyscan microscopy that provides a 1.4-fold increase in lateral resolution (∼ 140 nm) as compared with conventional confocal microscopes^23^. To compare the localization and dynamics of the *cis*-Golgi to the rest of the Golgi we crossed the *cis*-Golgi mCherry-MEMB12 or mRFP-SYP31 with the medial-Golgi marker NAG1-GFP and performed live cell airyscan acquisitions in root epidermal cells^24–26^. We first performed 5 min time-laps acquisitions and found *cis*-Golgi compartments moving together with the medial-Golgi, consistently with the conventional Golgi stack model (**Fig. 1a, c**; **videos 1 and 2**). However, we additionally observed *cis*-Golgi compartments moving independently from the medial-Golgi for both MEMB12 and SYP31 (**Fig. 1b, d**; **videos 1 and 2**). To be sure that the Golgi-independent compartments were completely separated from the Golgi in 3D, we acquired 3D-stacks by airyscan microscopy in living root epidermal cells. 3D-stacks were visualized and analyzed by IMARIS software to determine the 3D position of the Golgi-independent compartments from the medial-Golgi. Interestingly, we found *cis*-Golgi structures clearly separated in 3D from the medial-Golgi for both MEMB12 and SYP31, we also found Golgi-associated *cis*-Golgi (**Fig. 1e, f**; **video 3**; **Extended data 1e,f**). IMARIS surface rendering of reconstructed *cis*- and medial-Golgi objects additionally confirmed these observations (**Fig. 1g-n**; **video 3**; **Extended data 1g, h**). We next evaluated the proportion of the *cis* compartment that was independent from or associated with the medial-Golgi in our 3D acquisitions and found that Golgi-independent *cis* can reach up to more than half of the total population of *cis*-Golgi (**Fig. 1o**). These results challenge the traditional Golgi stack representation. Notably, we found a striking difference between the *cis*-markers: SYP31-positive compartments being mostly associated with the Golgi while MEMB12-compartments are mostly independent from it (**Fig. 1g-o**). These results suggest that the two *cis*-markers label two different populations of *cis*-Golgi compartments. Using IMARIS surface rendering we found that the *cis*-Golgi subpopulations are variable in size and display different morphologies (**Fig. 1g-n**). SYP31-compartments that associate with the rest of the Golgi are rounder and bigger than those labeled by MEMB12 which appear more fragmented (**Fig. 1g-n**, **Extended data 1e-h**). We next ranked the *cis*-compartments according to their distance to the rest of the Golgi. Interestingly, our results show that the largest *cis*-compartments are found in close proximity of the Golgi while the smallest are the most distant from the Golgi (**Fig. 1p**, **Extended data 1i**). We visualized this correlation for both MEMB12 and SYP31 *cis*-Golgi markers although the profiles were different between the two markers. We calculated that SYP31-compartmens are packed in one population that stayed within a 3-4 µm distance from the rest of the Golgi (**Fig. 1p**, **Extended data 1i**). Contrastingly, MEMB12-compartment are spread in a binomial distribution, one major population is located within a 3-4 µm distance from the rest of the Golgi, similar to SYP31, while a second population locates between 10-15 µm from the Golgi (**Fig. 1p**, **Extended data 1i**). These results suggest that the *cis*-compartments form larger structures in proximity of the Golgi and that at least two subpopulations of *cis*-Golgi co-exist, SYP31 that is close from the Golgi and MEMB12 that can be both close and distant from it. Thus, our next question was to address how these independent *cis*-Golgi behave dynamically. To address this question, we setup a workflow composed of lived airyscan microscopy acquisition, tracking of individual objects in both channels using the Detect Track and Co-localize (DTC) plugin of imageJ^27^. This plug-in identifies the interactions occurring between compartments of two channels and calculates their durations. Within our 5 min time-lapse acquisition, we found that the *cis*-Golgi compartments could either stay associated with the medial-Golgi (**Fig. 2a, d**; **videos 1 and 2**), remains independent from it (**Fig. 2b, e**; **videos 1 and 2**), or show transient interactions with the Golgi (**Fig. 2c, f**; **videos 1 and 2**; **Extended data 2a, e**). Importantly, in addition to the MEMB12-compartments we often and consistently detected a network of MEMB12-interconnected tubules that displays a super-fast kinetic and that is locally wrapped or packed into denser clustered structures that eventually associate with the medial-Golgi (white stars in **Fig. 1a, b, e**; **Fig. 2a-c**; **videos 1, 2, 4, 5**; **Extended data 2a**). Next, we quantified the proportion of MEMB12-structures that either associate with, are independent from, or show transient interaction with the Golgi. Our results showed clear differences between MEMB12 and SYP31 markers. MEMB12 mainly displayed transient interactions with the medial-Golgi, the rest of the MEMB12 population equally remains either associated to or independent from the Golgi (**Fig. 2g, h**). In contrast, SYP31 mainly remains associated to the medial-Golgi, or display transient interactions with it, while very few Golgi-independent entities were detected (**Fig. 2g, h**). If we now consider the population of transient interactions only, we quantified that the number of associations is equal to the number of dissociations within this population, for both MEMB12 and SYP31 markers (**Fig. 2i**). These results further argue that the interaction is only transient. Again, we noticed that MEMB12 displays about four times more association/dissociation than SYP31 (**Fig. 2i**). Our results are consistent with those we obtained in 3D IMARIS-reconstructed images (**Fig. 1g-o**). Next, we quantified the time of interaction and found out that the duration is around 15 seconds in average, for both MEMB12 and SYP31 markers (**Fig. 2j**). To be sure that the transient interactions we detected are not due to the loss of the objects during the tracking, we checked the average duration of one track and found that it was superior (around 22 sec) to the average association time (**Extended data 2a, b, e, f**), this gave us the upper edge of our quantification method. To obtain the lower edge, we evaluated the stochastic background of the method, in other words: what is the chance to detect an interaction just by chance? To check this, we either turned one channel by 180° or mirrored it vertically or horizontally and keep the other channel still. We found that the number of Golgi-associated *cis* is reduced close to 0 and that the number of Golgi-independent *cis* is much higher (**Extended data 2c, g**). Furthermore, the average interaction time is strongly reduced to 5-6 seconds (**Extended data 2d, h**). Thus, the interaction events we see are not due to random movements.

**Figure 1.**
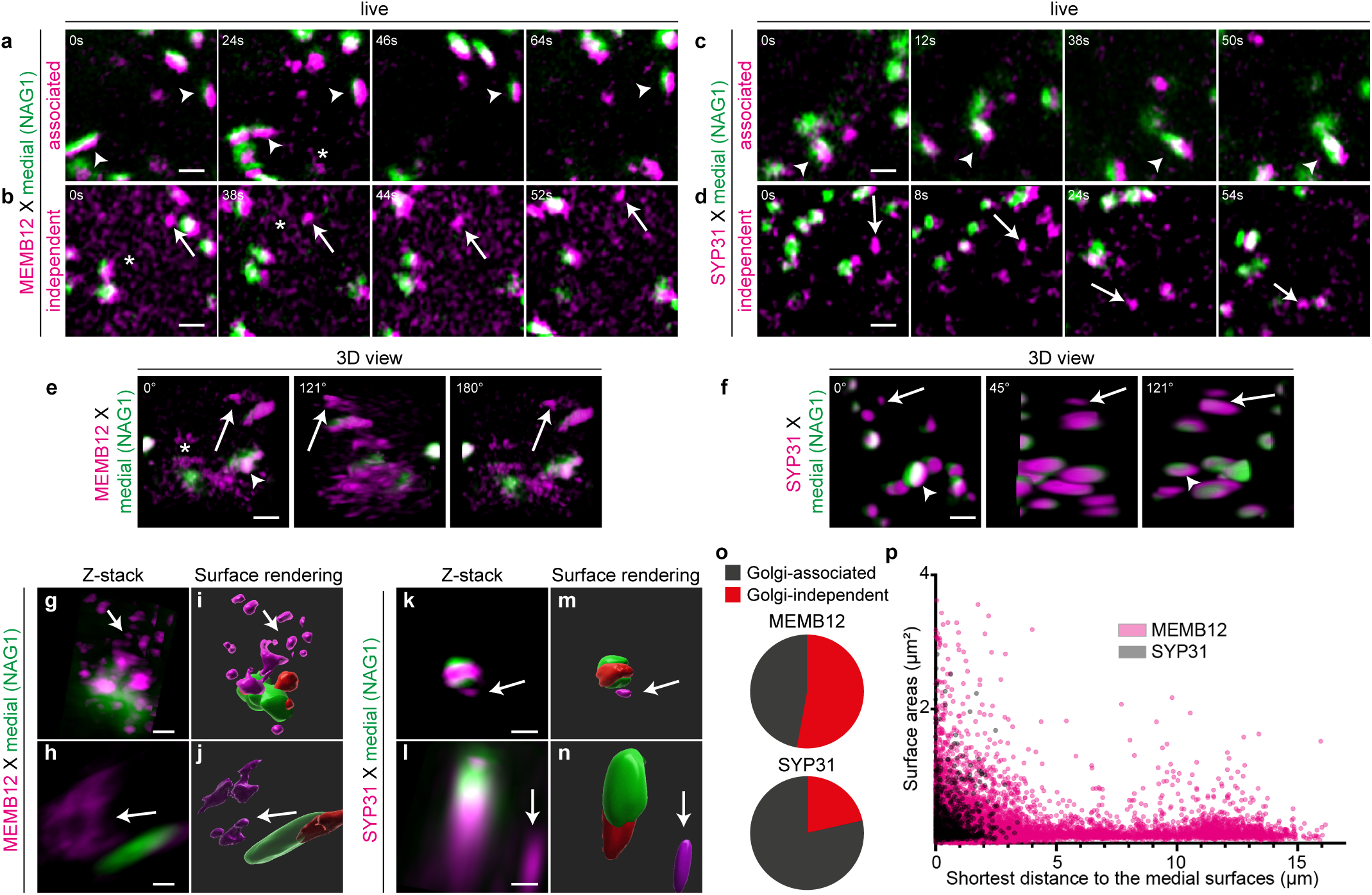
The cis-Golgi contains two subpopulations, the MEMB12-population is mostly Golgi-independent while the SYP31-population is mostly Golgi-associated. (**a-d**) Time-lapse airyscan acquisition in root epidermal cells of either mCherry-MEMB12 (cis-Golgi) x NAG1-EGFP (medial-Golgi) (**a, b**) or mRFP-SYP31 (cis-Golgi) x NAG1-EGFP (**c, d**) showing cis-compartments that remains either associated with (white arrowheads) or remains independent from the medial-Golgi (white arrows). White stars in a, b, e indicate MEMB12 tubular-like structures. (**e, f**) 3D airyscan acquisition of either mCherry-MEMB12 x NAG1-EGFP (**e**) or mRFP-SYP31 x NAG1-EGFP (**f**) showing that Golgi-independent compartments are not connected to other Golgi in the Z dimension. (**g-n**) 3D reconstruction (**g, h, k, l**) and surface modelling by the IMARIS software (**i, j, m, n**) of either mCherry-MEMB12 x NAG1-EGFP (**g-j**) or mRFP-SYP31 x NAG1-EGFP (**k-n**). (**o**) quantification of IMARIS surface modelling (n= 27 015 Golgi compartments identified out of 12 cells for MEMB12 x NAG1, n= 3 727 Golgi compartments identified out of 12 cells for SYP31 x NAG1) confirming that while SYP31 population is mostly associated to the Golgi, MEMB12 is mostly independent from the Golgi. (**p**) Ranking of IMARIS-modelled area (in µm²) of MEMB12- and SYP31-compartments according to their distance to the closer medial compartment. The MEMB12 population is more spread than SYP31, with one peak close to the Golgi (within 3-4 µm) and another peak 10-15 µm away from the Golgi. For both SYP31 and MEMB12, the closer to the Golgi and the largest are the compartments. All scale bars are 1µm.

**Figure 2.**
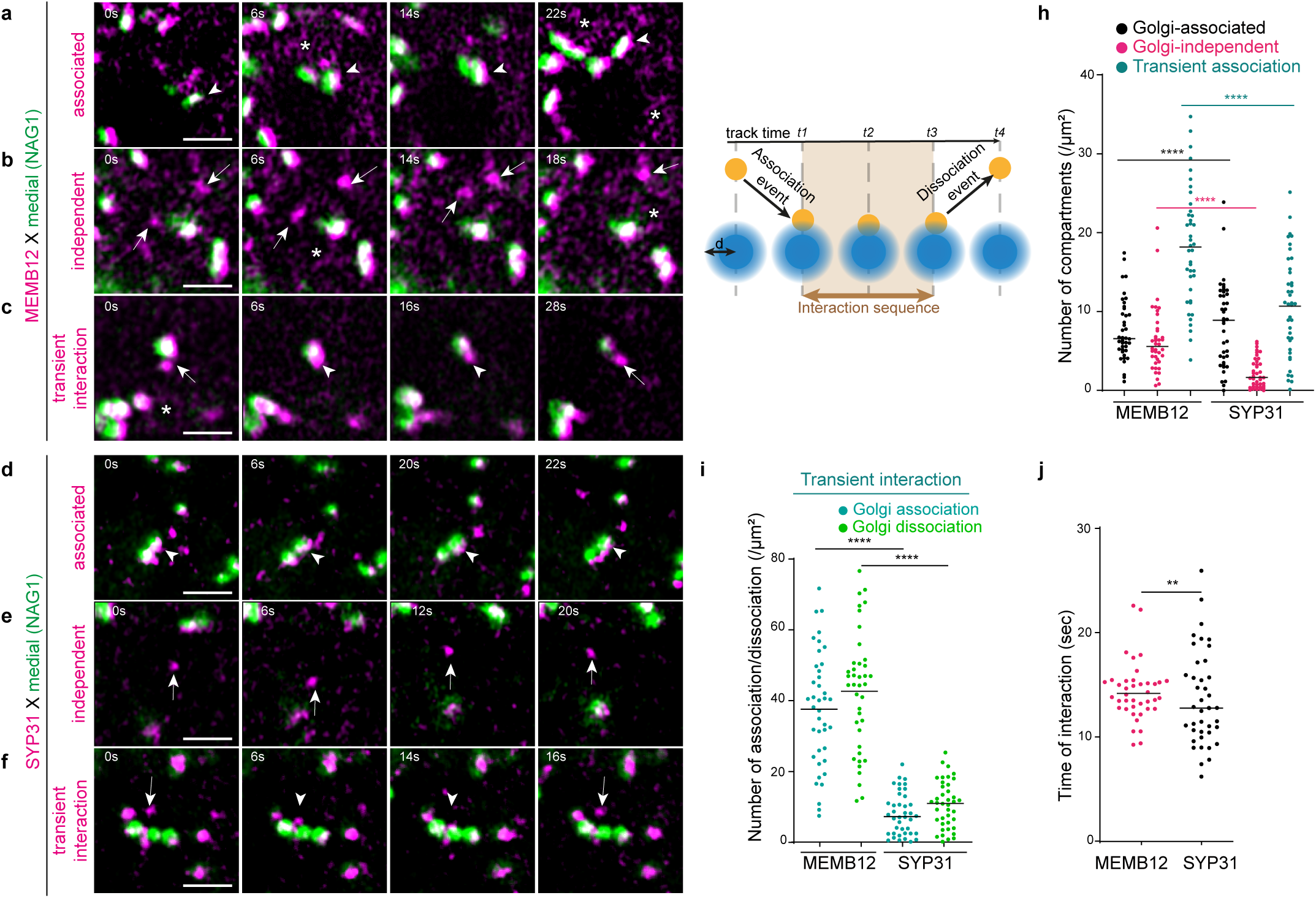
MEMB12 cis-Golgi is highly dynamic and displays transient interactions with the medial-Golgi. (**a-f**) Time-lapse airyscan acquisition in root epidermal cells of either mCherry-MEMB12 x NAG1-EGFP (medial-Golgi) (**a-c**) or mRFP-SYP31 x NAG1-EGFP (**d-f**). (**a, d**) Representative time-lapse of an association of either MEMB12- (**a**) or SYP31-compartments (**d**) with the medial-Golgi. (**b, e**) Representative time-lapse of Golgi-independent movements of either MEMB12- (**b**) or SYP31-compartments (**e**) away from the medial-Golgi. (**c, f**) Representative time-lapse of transient movements of either MEMB12- (**c**) or SYP31-compartments (**f**) with the medial-Golgi. White stars in **a, b, c** indicate MEMB12 tubular-like structures. (**g**) Schematic representation of the quantification workflow used for time-lapse acquisitions. Each compartment is followed for a certain time before it disappears from the field of acquisition, this is the track time. An association is detected if an opposite compartment crosses the association-zone (light blue) during the track time. The association is followed for a certain amount of time, this is the interaction sequence. (**h**) Number of MEMB12 or SYP31 compartments (normalized by cell area in µm²) that are either Golgi-associated, Golgi-independent or display a transient interaction with the Golgi. MEMB12 is more dynamic and independent from the Golgi than SYP31 (**** p<0.0001, by two-sided Wilcoxson and t-test rank sum tests). (**i**) Within the population of transient interaction, SYP31 displays less events of association and dissociation than MEMB12 (**** p<0.0001, by two-sided Wilcoxson and t-test (welch correction) rank sum tests). (**j**) Both MEMB12 and SYP31 have an interaction time around 15 sec with the medial-Golgi. n= 40 cells in all set of data ‘** p<0.0021, t-test rank sum test). All scale bars are 1µm.

### MEMB12 subpopulation dynamically associates with the ERES and is loaded with luminal cargos

To decipher whether the MEMB12*-*structures could be some ER-Golgi intermediate compartments we tested whether the MEMB12-compartments could associate with the ERES in plant cells and whether some cargos could be loaded into it. To check this possibility, we crossed the ERES marker MAIGO5 (MAG5)-GFP^28,29^ with the mCherry-MEMB12 fluorescent line and performed live cell airyscan time laps two-colors acquisitions in root epidermal cells (**video 6**). Interestingly, the MEMB12-population mainly displays either transient interactions or remains independent from the ERES in equal proportion (**Fig. 3a-d**; **video 6**). The number of MEMB12-structures that remained associated to the ERES all over the 5 min acquisition time is close to 0 further suggesting that MEMB12/ERES interactions are never stabilized, contrastingly to MEMB12/NAG1-medial Golgi interactions (**Fig. 3d, 2h**). Furthermore, within the dynamic population that display transient interactions, the number of association and dissociation is equal supporting the dynamic nature of the MEMB12-compartments regarding the ERES (**Fig. 3e**). We calculated the average interaction time and found that MEMB12-compartments stay associated with ERES for around 12 sec (**Fig. 3f**). Our upper edge control shows that the average track time is higher (around 22 sec) than the average association time (**Extended data 3a**). The lower edge 180° rotation and mirrored stochastic controls shows that the number of ERES-independent MEMB12-compartments are increased while the association time is decreased (**Extended data 3b, c**). Together, these results show that the *cis*-Golgi is able to be connected with the ERES in a dynamic way. As the ERES are supposed to be the sites of cargo loading, we established the Retention Using Selective Hook (RUSH)^4,30^ system in plants to test the possibility that some cargos, that are stacked at the ER in the first place and then synchronously released upon application of a chemical, are actually loaded into the MEMB12-compartments. The ER-hook we designed is composed from a ER-resident signal, the HDEL peptide^31^, a bait that is a mutated version of the FKBP-rapamycin binding (FRB) domain of the Human kinase TOR able to self-interact with itself^32^, and the fluorescent protein TagBFP2 (**Fig. 3g**). As a cargo we employed the fluorescent marker sec-EGFP that is widely used to identify ER-to-Golgi trafficking mechanisms in plants^33^, to which we fused a FKBP domain. We combined in one construct the ER-hook and the secretory cargo with the self-cleaving peptide P2A in between them. As the affinity of the mutated FKBP is higher for FKBP-ligand than for itself, the addition of FKBP-ligand will release the cargo construct from the ER (**Fig. 3h**). In transient expression in the *Arabidopsis* cotyledons, we could perfectly see an ER-pattern for sec-EGFP-RUSH that perfectly matches the ER-pattern of the HDEL-hook (**Fig. 3i-k**). In stable expression in the Arabidopsis root epidermis, the fluorescence of the HDEL-hook was too weak to be detected but sec-EGFP-RUSH was efficiently stacked at the ER network (**Extended data 3d**). Upon cargo release by the FKBP-ligand, we observed that sec-EGFP-RUSH labels some endomembrane compartments that either closely associate or co-localize with the MEMB12-compartments, both in transient expression in cotyledons (**Fig. 3 l-n**) and stable expression in roots (**Fig. 3o-x**). We also noticed that the sec-EGFP-RUSH is reaching pro-vacuole-like structures within 10 min and larger vacuoles within 60-80 min after cargo release both in roots (**Extended data 3e-g**) and cotyledons (**Extended data 3h-j**). To quantify the trafficking of sec-EGFP-RUSH through the Golgi, we performed airyscan localization analyses of sec-EGFP-RUSH at different time points after cargo release, up to 20 min, in the root system. Interestingly, we found that the co-localization of sec-EGFP-RUSH with the mCherry-MEMB12-compartments is strongly increased after 5 min of cargo release and progressively decreases already from 10 min after release (**Fig. 3y**). These results show that the luminal sec-EGFP-RUSH is indeed crossing the MEMB12 subpopulation at early time points after leaving the ER.

**Figure 3.**
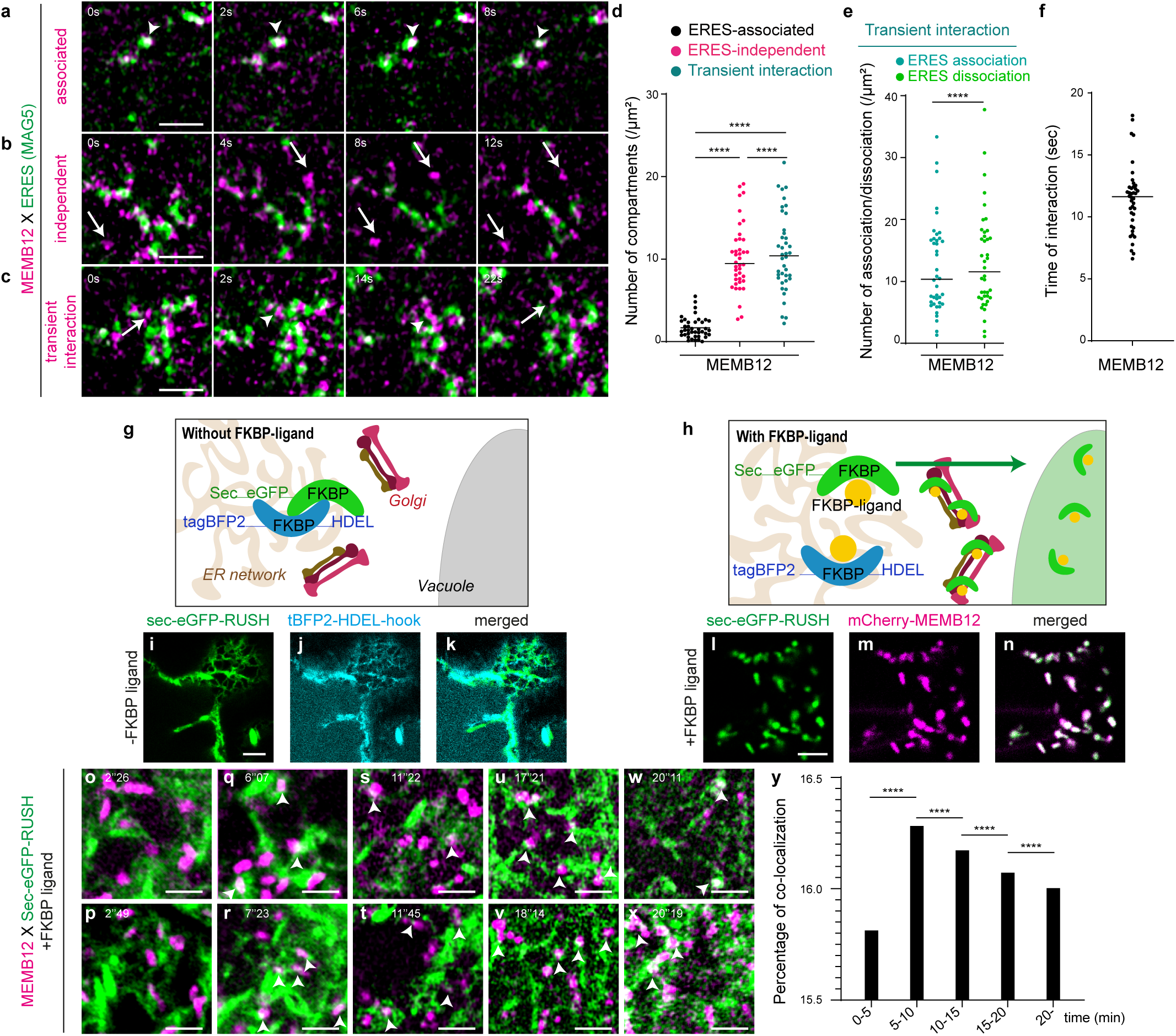
MEMB12 cis-Golgi displays transient interactions with the ERES and is crossed by the luminal cargo sec-GFP early after inducible-release from the ER. (**a-c**) Time-lapse airyscan acquisition in root epidermal cells of mCherry-MEMB12 x MAG5-GFP (ERES). (**a**) Representative time-lapse of MEMB12-compartments either associated to the ERES, (**b**) independent from the ERES or (**c**) in transient interaction with the ERES. (**d**) Number of MEMB12-compartments (normalized by cell area in µm²) that are either ERES-associated, ERES-independent or display a transient interaction with the ERES (n= 40 cells, **** p<0.0001, by two-sided Wilcoxson and t-test (welch correction) rank sum tests). (**e**) Within the population of transient interaction, MEMB12 displays equal number of association and dissociation events with the ERES (n= 40 cells, **** p<0.0001, by two-sided Wilcoxson rank sum test). (**f**) MEMB12 has an interaction time around 12 sec with the ERES (n= 40 cells). (**g, h**) Schema of the sec-EGFP-RUSH retained at the ER by the HDEL-hook by FKBP-dimerization and released upon incubation with FKBP-ligand. (**i-n**) Confocal images of transient transformation in epidermal cotyledon cells of the sec-EGFP-RUSH construct. (**i-k**) The sec-EGFP-RUSH (**i**) is retained by the tagBFP2-HDEL-hook at the ER-network (**j**, merged in **k**). (**l-n**) Shortly after release, sec-EGFP-RUSH (**l**) strongly co-localizes with mCherry-MEMB12 (**m**) as visible in the merged picture (**n**). (**o-x**) Airyscan acquisition of root epidermal cells stably expressing mCherry-MEMB12 together with sec-EGFP-RUSH. The timing of incubation with the FKBP-ligand is indicated in the upper-left corner of each images. (**y**) Quantification of the percentage of co-localization between sec-EGFP-RUSH and mCherry-MEMB12 up to 20 minutes of incubation with the FKBP-ligand. n=40 cells for each experiment in **d-f**, n= 548 cells out of 36 roots in **y** (**** p<0.0001 by two-sided Wilcoxon rank sum test). Scale bars are 1 µm in **a-c**, 4 µm in **h-j** and 2 µm in **o-x**.

### The MEMB12 subpopulation is a tubulo-vesiculated network mostly independent from the Golgi but displaying local Golgi-cisterna association

In our airyscan time-lapse acquisitions, we not only found dynamic MEMB12-compartment but also a network of super-dynamic MEMB12-interconnected tubules (white stars in **Fig. 1a, b, e**; **Fig. 2a-c**; **videos 1, 2, 4, 5**; **Extended data 2a**). However, airyscan microscopy has around 140 nm resolution in lateral dimension and might not have sufficient resolution to fully distinguish membrane tubules. To confirm that MEMB12 labels such as network we employed a super-resolution microscopy approach using STED microscopy^34,35^. To obtain the best resolution we employed the τ-STED technology that uses Fluorescence Lifetime Imaging Microscopy (FLIM) approach to discriminate the background from the fluorophores^36,37^. Additionally, we used nanobodies, the smallest known antibodies, coupled to STED-compatible fluorophores to obtain the maximal resolution^38,39^. In our images, we calculated that the lateral resolution is 63 nm (+/- 12 nm) in τ-STED while we obtained 166 nm (+/- 19 nm) in airyscan microscopy (see methods section). We performed two-colors τ-STED microscopy with the mCherry-MEMB12 marker and the medial-Golgi marker NAG1-GFP. Strikingly, the localization pattern of the two markers appeared completely different. The medial-Golgi is a cisterna that looks either flat, like a rice grain, or circular with an edge localization of NAG1-GFP, like a donut, depending on the orientation of the cisternae (**Fig. 4a, d, g**). Contrastingly, the MEMB12-compartments were a tubulo-vesicular network with small vesicles connected by tubules, similar to the VTC/ERGIC of animal cells (**Fig. 4b, e, h**). To quantify these observations, we used a morphometric workflow that calculates three parameters: the area, the circularity and the solidity (the circularity measures if a structure is carved or branched) of the structures. We found that the area of MEMB12-structures was higher than for NAG1 but the values were also much more spread (**Fig. 4j**). The circularity of the medial-Golgi was seemingly spread in two groups, we found out that this was due to the orientation of the cisterna, either like a rice grain or like a donut (**Fig. 4f**, **Extended data 4a**). The circularity and solidity values of MEMB12-structures were lower than NAG1 but again more spread than for NAG1 (**Fig. 4j**, **Extended data 4a**). These results indicate that MEMB12-structures are more heterogeneous in shape than NAG1-medial-Golgi and quantitatively confirm that the MEMB12 compartments are not like Golgi-cisternae but are rather a mix of branched tubular network and vesicles (**Fig. 4a-i**). While the MEMB12 network appears mostly independent from the medial-Golgi cisterna, we consistently see close associations or intertwined connection of the MEMB12 compartments with the medial-Golgi (**Fig. 4c, f, i**). In proximity of the medial-Golgi we detected either several vesicle-shaped structures or more expanded structures (**Fig. 4f, i**), consistently to what we described earlier by IMARIS (**Fig. 1g-j**) or what was described by electron microscopy at the vicinity of the Golgi stack^7^. The mCherry-MEMB12 construct is under the control of the mild expressing promoter pUBQ10. Thus, we wanted to address whether the network we observed is an artefact due to the mild expression of the protein or whether we could also observe this network in native condition. For this, we performed an immuno-localization of MEMB12 protein with an antibody recognizing specifically MEMB12 and its close homolog MEMB11^40^. Our τ-STED results indicate that we could detect a similar tubulo-vesicular network, mostly independent from the medial-Golgi with local association or imbrication with the medial-cisterna (**Fig. 4k-s**). We noted that sometimes MEMB12 was labeling structures that appear like a tubulo-vesicular cluster independent form the Golgi or like a membrane sheet or cisterna associated with the medial-Golgi (**Fig. 4p, s**). The morphometric analysis of anti-MEMB12 basically show similar results as for the mCherry-MEMB12 with spread values for the area, circularity and solidity representative of a mix between tubules and vesicles (**Fig. 4t**). Hence, we confirmed that the tubulo-vesicular network labeled by MEMB12 is present in plants that do not express any *cis*-Golgi fluorescent markers. Strikingly, we performed τ-STED on p35S::mRFP-SYP31 and saw a cisterna-like structure, similar to the medial-Golgi although being smaller than NAG1-cisterna (**Extended data 4c-m**). Cisterna-like structures were also observed for pUBQ10::YFP-SYP32 or pSYP32::GFP-SYP32 Arabidopsis lines (**Extended data 4c-k**) and were also smaller than the medial-Golgi (**Extended data 4l, m**, area). These results are consistent with the IMARIS morphometric analyses where SYP31 compartments appeared bigger and less fragmented than MEMB12 compartments (**Fig. 1g-j**). Finally, to confirm our findings by yet another independent approach than τ-STED processing, we performed Expansion Microscopy (ExM) to physically expand our samples by a 4X factor and acquire 2D images and 3D-stacks by conventional confocal microscopy. In our ExM images we obtained a lateral resolution of 91 nm (+/- 8 nm). Our results clearly show that the MEMB12-labeling is distributed over a tubulo-vesiculated network that is mostly independent from the medial-Golgi (**Fig. 4u-w**) but that can locally display contact with it (**Fig. 4x-ac**). Contrastingly, the medial-Golgi is a cisterna, with an accumulation of NAG1-EGFP at the edges of the cisterna, like what we described in τ-STED (**Fig. 4u, x, aa**; **Fig. 4a, d, g, k, n, q**). Furthermore, we acquired 3D ExM stacks, reconstructed them using IMARIS surface rendering and confirmed that MEMB12 appears like a Golgi-independent fragmented network while NAG1 is more an isotropic compartment (in **Fig. 4ad-ag**, **video 7**, white arrows). Interestingly, some part of the MEMB12 network locally surrounds or entangle the medial-Golgi (**Fig. 4ad-ag**, white arrowheads) and tubular structures seems to connect the Golgi-independent and Golgi-associated MEMB12 network (**Fig. 4ad-ag**, white stars). Altogether, our τ-STED and ExM results demonstrate that the morphology of MEMB12-structures is a Golgi-independent tubulo-vesiculated network that can intertwine the Golgi cisternae.

**Figure 4.**
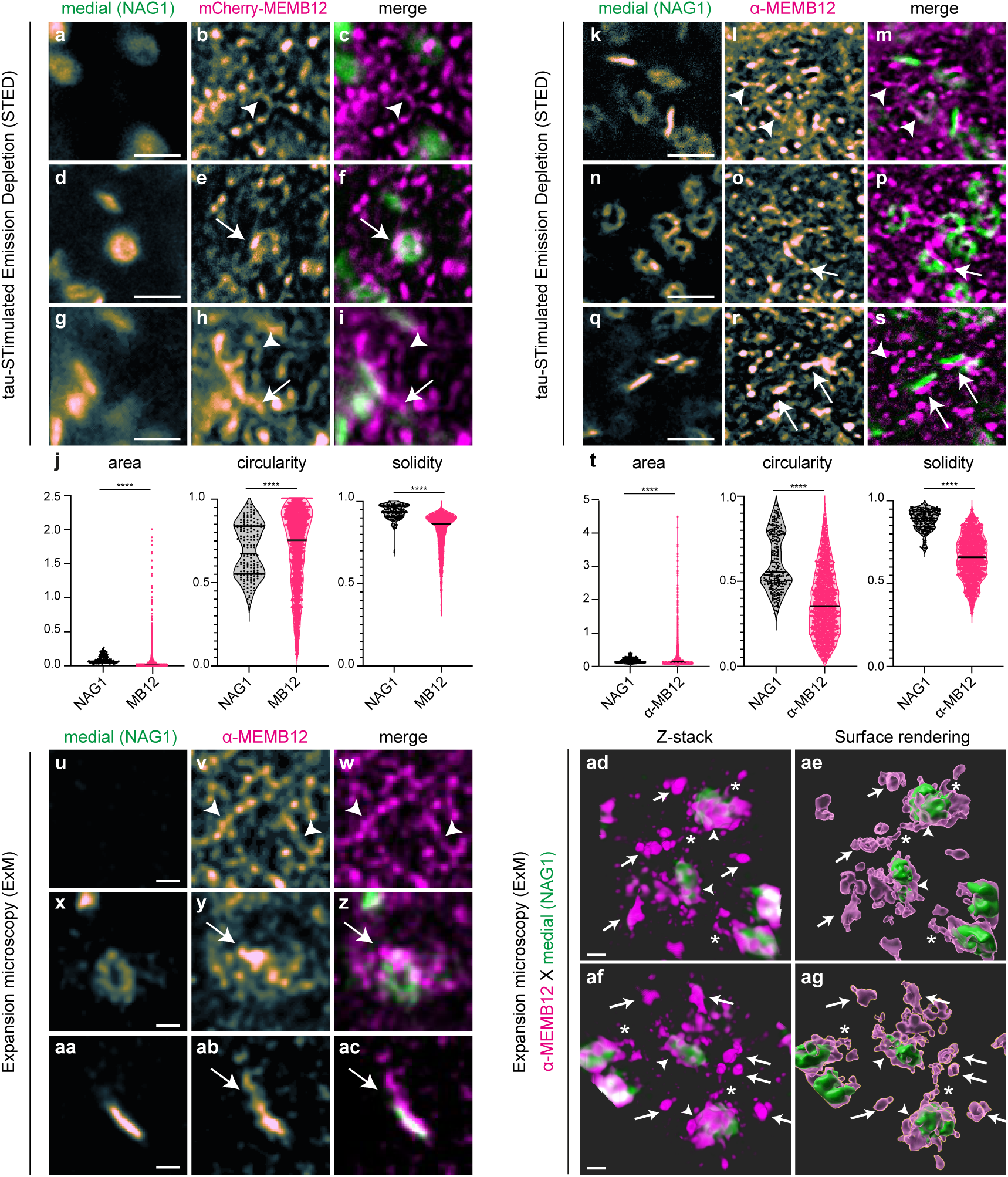
MEMB12 cis-Golgi is a tubular network mostly independent from the Golgi that displays local intertwined connection with the medial-Golgi. (**a-t**) Lifetime τ-Stimulated Emission Depletion (τ-STED) of either mCherry-MEMB12 x NAG1-EGFP (**a-i**) or α-MEMB12 immuno-stained compartments in NAG1-EGFP expressing plants (**k-s**). (**j, t**) Morphometric quantification of **a-i** and **k-s**, respectively. The circularity of NAG1 was split in two groups, the highest one corresponds to donut-like structures while the lowest group corresponds to rice grain-like structures (see Extended data Fig. 4). The solidity of NAG1 was high indicative of a homogenous and quiet isotropic structure. Contrastingly, the circularity and solidity of MEMB12 were a spread population with lower values for anti-MEMB12 than for mCherry-MEMB12 indicative that anti-MEMB12 is better revealing the tubular network than mCherry-MEMB12. n=3049 compartments for mCherry-MEMB12, n=161 for NAG1-EGFP i n **j**; n=1837 compartments for α-MEMB12, n=238 for NAG1-EGFP in **t**. (**u-ag**) 4X Expansion Microscopy (ExM) performed on α-MEMB12 immuno-stained compartments in NAG1-EGFP expressing plants. (**u-ac**) 2D acquisitions, (**ad-ag**) 3D acquisitions, reconstruction and surface rendering by IMARIS. For both τ-STED and ExM, MEMB12 is localized on a tubulo-vesicular network (white arrows and arrowheads) while NAG1 is a cisterna looking a rice grain or a donut, depending on its orientation (note the edge localization of NAG1). The MEMB12 tubulo-vesicular network is mostly independent from the medial-Golgi (arrowheads in **b, c, l, m, v, w, ad-ag**) but intertwined connection are often observed between MEMB12-network and NAG1-cisterna, like on the white arrows in **e, f, h, i, o, p, r, s, y, z, ad-ag**. The white stars in **ad-ag** indicate tubular connections between Golgi-associated and Golgi-independent MEMB12-network. Connections between MEMB12 and NAG1 can further extend to cisterna-like compartments labelled by MEMB12 that lies close to the medial-cisterna (arrows in **r, s, ab, ac**). * p<0.05; ** p<0.005; *** p<0.0005; **** p<0.0001 by Wilcoxon rank sum test. Scale bars are 1µm in **a-i**, **k-s** and 250 nm in **u-ag**.

### The dynamic behavior of MEMB12 subpopulation is regulated by sphingolipid membrane composition and its contacts with the Golgi cisternae

MEMB12 is localized on a tubular network. Interestingly, it is known that the sphingolipids ceramides containing Very-Long Chain Fatty Acids (VLCFAs) of 24 atoms of carbon (C24-ceramides) favor the formation of tubular structures through the formation of interdigitated phases^41^. Thus, we used the FumonisinB1 (FB1), an inhibitor of the Arabidopsis ceramide synthases LOH1 and LOH3 that specifically synthesize C24-ceramides^42^. As an additional tool, we used Metazachlor (Mz), that specifically inhibits the synthesis of C24 fatty acids that will next be integrated into sphingolipids^43,44^. However, FB1 is more specific towards ceramides but generates an accumulation of free Long Chain Bases (LCB) while Mz is not specific towards a sphingolipid specie but has less side-effects^42^. Our results show that FB1 treatment, but not Mz, strongly reduces the number of MEMB12-compartments as well as the number of NAG1 medial-Golgi (**Fig. 5a, b, d, e**; **Extended data 5a**). These results suggest that FB1 inhibits the *de novo* generation of MEMB12-compartments and Golgi stacks. As a result of the loss of MEMB12 and medial-Golgi upon FB1, the number of Golgi-associated, Golgi-independent and transient association of MEMB12 with the medial-Golgi is strongly reduced upon FB1, but not upon Mz (**Extended data 5b, c**). If we now only look at the population that transiently associate with the medial-Golgi, we quantified that the number of associations/dissociations was strongly reduced upon both FB1 or Mz (**Fig. 5c, f**). These results suggest that the effect of Mz is milder as compared to FB1 and the dynamic interaction of MEMB12-compartments with the medial-Golgi is dependent on the acyl-chain length composition of sphingolipids. As controls, the velocity of MEMB12-compartments or the time of association with the medial-Golgi was not altered neither upon FB1 nor upon Mz (**Extended data 5d-g**). Additionally, to these dynamic quantifications, we noticed in our airyscan time-lapse acquisitions that FB1 or Mz treatment alters the MEMB12 tubular-network, some tubules are still visible in the surroundings of the Golgi but the main tubular-network is absent in Golgi-free zones (**videos 8 and 9**). These results suggest that sphingolipids are involved in the generation of the tubular-network and its dynamics and that the interaction with the Golgi could stabilize the MEMB12-network, as we already suggested in several set of experiments earlier in this study. Thus, next we addressed whether the contact of the MEMB12-compartments with the rest of the Golgi additionally contributes to its dynamics. Our dynamic airyscan, IMARIS and τ-STED results suggest that SYP31 is mostly a stable cisterna close to the medial-Golgi while MEMB12 is mostly a highly dynamic tubulo-vesiculated structure that can sometimes be stabilized at the Golgi cisternae. Thus, we wondered whether the MEMB12-network displays dynamic association with SYP31-cisterna. To address this, we crossed mCherry-MEMB12 into EGFP-SYP31 and performed airyscan time laps acquisitions. Indeed, we quantified that MEMB12 is mostly transiently associated with SYP31, with an average time of association between 12-15 sec, consistently to what we observed for the medial-Golgi (**Fig. 5g-j**; **video 10**). Our controls show that the track time is higher than the association time and that the rotation or inversion of one channel results in the increase of the number of SYP31-independent MEMB12-compartments and decrease of the association time (**Extended data 6a-c**). Our IMARIS analyses showed that the MEMB12-compartments are getting enlarged when they get close to the medial-Golgi, suggesting that they get stabilized at the Golgi (**Fig. 1g-j**). Thus, we quantified the velocity of MEMB12-compartments that are either associated with or independent from the Golgi and found that the speed of MEMB12-compartments was lower when it is associated with either SYP31-*cis* or NAG1-medial Golgi (**Fig. 5k, l**). Similarly, the SYP31 compartments that are independent from the medial-Golgi are faster than the ones associated with it (**Extended data 6d**). Altogether, our dynamic and morphometric analysis suggest that the MEMB12-stucture is mostly dynamics but gets stabilized and enlarges into bigger structures when it locally contacts the *cis*-Golgi cisterna. Indeed, in our videos we often and consistently see that the dynamic MEMB12-network gets locally wrapped into a denser clustered structure at the proximity of either NAG1-medial or SYP31-*cis* cisternae (**videos 1, 2, 4, 5, 10**).

**Figure 5.**
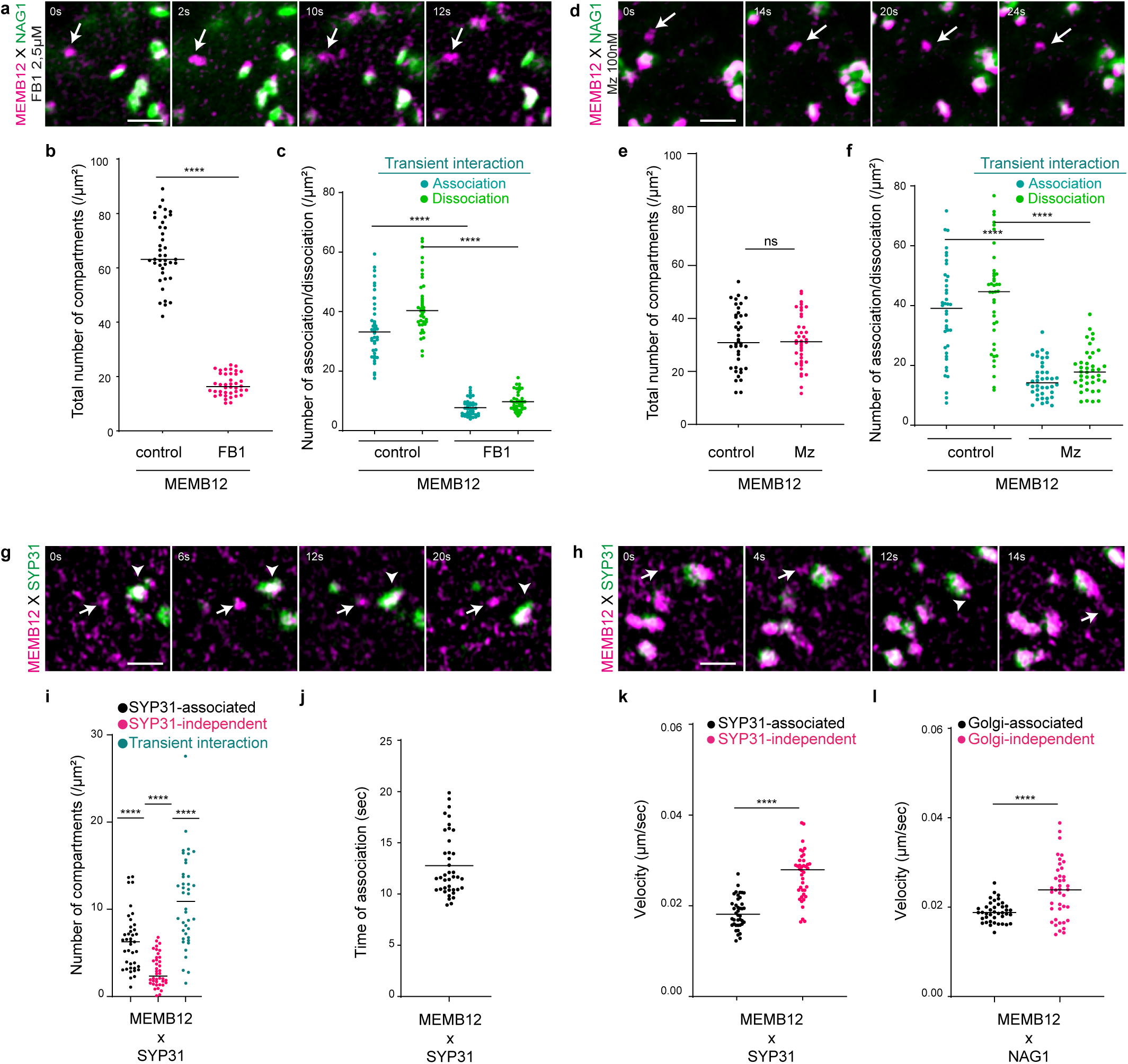
The dynamic behaviour of MEMB12-structures is regulated by sphingolipids and its interaction with the Golgi-cisternae. (**a-f**) Root epidermal cells expressing mCherry-MEMB12 x NAG1-EGFP (medial-Golgi) upon either 2.5µM FB1 (**a-c**) or 100 nM Mz (**d-f**) treatment. (**a, d**) Time-lapse airyscan acquisitions showing Golgi-independent MEMB12 compartments. (**b, e**) Quantification of the total number of MEMB12-compartments showing a severe decrease upon FB1 but not upon Mz (n=40 cells for either FB1 or Mz, **** p<0.0001 by t wo-sided Wilcoxon’s rank sum test for FB1 and two-sided t-test rank sum test for Mz). (**c, f**) MEMB12 displays less association and dissociation with the medial-Golgi upon either FB1 (**c**) or Mz (**f**) treatment than control condition (n=40 cells for either FB1 or Mz, **** p<0.0001 by Wilcoxon’s rank sum test for FB1 and t-test’s (welch correction) rank sum test for Mz). (**g-l**) Root epidermal cells expressing mCherry-MEMB12 x EGFP-SYP31. (**g, h**) Time-lapse airyscan acquisition of mCherry-MEMB12 x GFP-SYP31 showing independent and associated movement of MEMB12 with SYP31 (**g**) or transient interaction with SYP31(**h**). (**i-l**) Quantification of the MEMB12 dynamics, relative to SYP31. (**i**) MEMB12 is mostly displaying transient interaction or stable association with SYP31 (n= 40 cells, **** p<0.0001 by Kruskal-Wallis test and two-sided Wilcoxon’s rank sum tests). (**j**) The interaction time of MEMB12-compartments with SYP31 is close to 12 seconds. (**k, l**) The velocity of MEMB12-compartments is decreased when it associates to either SYP31 (**k**) or the medial-Golgi (**l**) (n= 40 cells, **** p<0.0001 by two-sided t-test (welch correction) rank sum test). All scale bar are 1µm.

## Conclusion

Our study revises and transforms our understanding of the architecture and dynamics of the Golgi in a Eukaryotic multicellular system. Our results demonstrate that the *cis*-Golgi of Arabidopsis is constituted from different subpopulations, one is labeled by Qa-SNARE SYP31/32 and is mostly a stable cisterna-like compartment attached to the medial-Golgi. Another subpopulation is labeled by the Qb-SNARE MEMB12 and is highly dynamic network of membrane tubules and vesicles that is mainly independent from either the Golgi or the ERES but can still interacts with them either transiently or more stably. Why was this *cis*-Golgi network never observed before in plant cells? Actually, membrane tubules were suggested to exist between the ER and the plant Golgi for a long time^12,13,45^. However, previous immuno-EM experiments using *cis*-Golgi markers were all focused on the Golgi stack and, consistently with our results, all show a pearled membrane structure at the *cis*-side of the stack^7,40,46,47^. So up to now our understanding of the Golgi apparatus was biased by a stack-centered view of the Golgi and the governing idea that the *cis*-Golgi is a cisterna moving with the rest of the Golgi. GECCO was originally described using the SYP31 *cis*-Golgi marker^17,48^ which mostly labels a Golgi-associated cisterna in this study. We now propose that the highly dynamic Golgi-independent *cis*-Golgi MEMB12-tubulo-vesicular network is part of the ERGIC in plants and constitutes an early station of the GECCO, while the more stable Golgi-associated *cis*-Golgi SYP31-cisterna would correspond to a more mature part of GECCO. We show here that the MEMB12-tubular network dynamically interacts with both the ERES and the Golgi and is able to transport cargos from the ER to the Golgi due to that the sec-EGFP-RUSH construct is crossing the MEMB12-compartment shortly after induced ER-release. We noticed that the vacuole is also stained quickly after ER-release, thus we cannot exclude that luminal cargo follow an additional direct path from the ER as was described earlier for other type of cargos^49^. Our results show that the MEMB12-network is very dynamic in a lipid-dependent manner. More precisely, we found that the acyl-chain length of sphingolipids is involved in the generation of the MEMB12-tubular-network and in the dynamic interaction of MEMB12-compartments with the Golgi. Acyl-chain length of sphingolipids induces interdigitated membrane phase and the formation of tubules^41^. Thus, we postulate that a fine-tuned balance in membrane sphingolipid composition participates, probably in combination with other factors, to keep a flexible and dynamic MEMB12-tubular network. Consistently, our results show that the MEMB12-compartments are less tubular, enlarged and more stable when they get at the proximity of a Golgi-cisterna. Alternatively, acyl-chain length of sphingolipids could directly be involved in differential protein sorting, similarly to what has been described in yeast at ERES^50^. The results presented in this study open the way to the establishment of a revised model of the Golgi in which at least a part of the ER-to-Golgi trafficking occurs *via* a *cis*-Golgi intermediate tubular network. In this model, some discrete regions of the super-dynamic tubular MEMB12-network stabilize into an intertwined membrane sheet, eventually giving rise to a new *cis*-Golgi cisterna. This could explain how the Golgi maintains its integrity while being constantly crossed by cargos, by forming new *cis*-cisternae from the intermediate network. In mammalian cells, the ERGIC markers ERGIC-53 and Rab1 localize not only to the structures that are apart from the Golgi ribbon but also to the *cis*-face of the main Golgi body, suggesting that the similar organization of the intermediate compartments at the ER-Golgi interface is shared across kingdoms^5,51,52^. Our work paves the way to future exploration on this topic, more specifically on how the Golgi is generated and what is the role of membrane trafficking and remodeling during cell responses to environmental stimuli and/or developmental cell-reprograming membrane processes.

## Acknowledgments

We are grateful to Prof. Ikuko Hara-Nishimura and Dr. Haruko Ueda for sharing published material. Imaging was performed at the Bordeaux Imaging Center (BIC), part of the National Infrastructure France-BioImaging supported by the French National Research Agency (ANR-10-INSB-04). We are grateful to the BIC team for their expertise, support and access to super-resolution microscopes as well as their help in quantification. We thank Dr. Yvon Jaillais and Dr. Patrick Moreau for the critical reading of the manuscript and helpful comments. This work was supported by French National Research Agency grants ANR CALIPSO (ANR-18-CE13-0025) and ANR FATROOT (ANR-21-CE13-0019) to Y.B., a PhD fellowship from the French government and distributed through the doctorate school in life and health sciences of Bordeaux to L.F.

## Author contribution statement

L.F performed all experiments in Fig. 1, 2, 3, 5 and Extended data 1, 2, 3, 5, 6. M.G established the τ-STED and ExM in Arabidopsis and performed all experiments in Fig. 4 and Extended data 4. P.L and M.M participated in the generation and selection of plant genetic material. F.C created the DTC plugin and helped in the quantification of time-lapse acquisition as well as τ-STED quantification. M.F-M participated with M.G to establish the ExM in Arabidopsis. C.P participated with M.G to establish the τ-STED microscopy in Arabidopsis. T.U provided unpublished SYP31 and SYP32 fluorescent marker line and provided scientific input. A.N provided scientific input and initial materials to this study. Y.I setup the initial airyscan live acquisition and quantification^27^, performed the cloning and solved the initial issues of the RUSH system in Arabidopsis and conceptualized the research. Y.B conceptualized and designed the research, got the financial support for, supervised all aspects of this study and wrote the manuscript. The figures were made by L.F and M.G. All authors read and provided inputs on the manuscript.

## Materials and correspondence

The materials and tools published in this study will be made available on demand to

## Data availability statement

All data generated or analysed during this study are included in this published article (and its supplementary information files).

## Legends of the videos

**Video 1. Airyscan time-lapse acquisition of MEMB12-mCherry and NAG1-EGFP in root epidermal cells.** MEMB12 is labelling vesicle-like structures as well as a tubular network (white star) that is highly dynamic. MEMB12-structures display both independent movements from the medial-Golgi (white arrow) and Golgi-associated movements. Scale bar is 1 µm, the time in seconds is indicated at the upper right.

**Video 2. Airyscan time-lapse acquisition of SYP31-mRFP and NAG1-EGFP in root epidermal cells.** SYP31 is mostly labelling vesicle-like structures that display Golgi-associated movements. Scale bar is 1 µm, the time in seconds is indicated at the upper right.

**Video 3. 3D-airyscan acquisition and IMARIS surface modelling of *cis*-/medial-Golgi compartments.** (**a**, **b**) 3D-airyscan images and surface modelling of either MEMB12-mCherry or SYP31-mRFP together with NAG1-EGFP in root epidermal cells. MEMB12-structures are more fragmented and dispersed than SYP31 that is closer to the Golgi and labelling the edge of a cisterna, like a donut. (**c**-**f**) Additional examples of 3D-reconstructed MEMB12 and SYP31 structures.

**Video 4. Single channel airyscan time-lapse acquisition of MEMB12-mCherry in root epidermal cells.** MEMB12 labels a highly dynamic tubular network that is locally wrapped or packed into denser clustered structures. Scale bar is 1 µm, the time in seconds is indicated at the upper right.

**Video 5. Another single channel airyscan time-lapse acquisition of MEMB12-mCherry in root epidermal cells.** MEMB12 labels a highly dynamic tubular network that is locally wrapped or packed into denser clustered structures. Scale bar is 1 µm, the time in seconds is indicated at the upper right.

**Video 6. Airyscan time-lapse acquisition of MEMB12-mCherry and MAIGO5-EGFP in root epidermal cells.** MEMB12-structures transiently associate (white arrows) with ERES structures labelled by MAIGO5-EGFP. Scale bar is 1 µm, the time in seconds is indicated at the upper right.

**Video 7. 3D-expansion microscopy (ExM) and IMARIS surface modelling of the MEMB12-network.** 3D-airyscan ExM images and surface modelling of MEMB12-mCherry together with NAG1-EGFP in root epidermal cells. The Golgi-independent MEMB12-network display intertwined connection with the medial-Golgi and is often found to surround it.

**Video 8. Airyscan time-lapse acquisition of MEMB12-mCherry and NAG1-EGFP in root epidermal cells treated with 100 nM Mz.** The Golgi-independent MEMB12-network disappeared upon Mz, only some part of the MEMB12-network remains close to the Golgi (white arrow). Scale bar is 1 µm, the time in seconds is indicated at the upper right.

**Video 9. Airyscan time-lapse acquisition of MEMB12-mCherry and NAG1-EGFP in root epidermal cells treated with 2.5µM FB1.** The Golgi-independent MEMB12-network disappeared upon FB1, only some part of the MEMB12-network remains close to the Golgi (white arrow). Scale bar is 1 µm, the time in seconds is indicated at the upper right.

**Video 10. Airyscan time-lapse acquisition of MEMB12-mCherry and SYP31-EGFP in root epidermal cells.** MEMB12 is labelling vesicle-like structures as well as a tubular network (white star) that is highly dynamic. MEMB12-structures display both independent movements from SYP31-compartments (white arrow) and SYP31-associated movements. MEMB12 labels a highly dynamic tubular network that is locally wrapped or packed into denser clustered structures close to SYP31-compartments. Scale bar is 1 µm, the time in seconds is indicated at the upper right.

## Methods

### Cloning, plant material and growth conditions

The following Arabidopsis transgenic fluorescent marker lines were used: pUBQ10:: mCherry-MEMB12 (Wave 127R)^24^, p35S::NAG1-EGFP^26^, p35s::GFP-SYP31, p35s::mRFP-SYP31, pSYP32::GFP-SYP32, pMAG5::MAG5-GFP^28,29^. To generate *p35S::GFP-SYP31* and *p35S::mRFP-SYP31*, the DNA fragments coding *p35S::XFP-SYP31-Nos3’* were amplified by PCR from the SYP31 vectors previously published^53^. They were cloned into pENTR/D-TOPO (Thermo Fisher Scientific/Invitrogen), and recombined into pKGW by LR Clonase II (Thermo Fisher Scientific/Invitrogen). To generate *pSYP32::GFP-SYP32*, the *Arabidopsis* genomic *SYP32* DNA fragment which contains 2.5 kb 5’ and 0.8 kb 3’ flanking sequences was amplified by PCR and cloned into pENTR/D-TOPO. The DNA fragment coding GFP was inserted at the 5’ side of SYP32 by In-Fusion HD Cloning Kit (Takara), and the whole sequence was recombined into pGWB1 by LR Clonase II. To generate *p35S::ST-mRFP*, first the DNA fragment coding ST was amplified by PCR and cloned into pENTR/D-TOPO, and recombined into pH7RWG2. Next, the DNA coding ST-mRFP was amplified from it and cloned into pENTR/D-TOPO again, and recombined into pGWB2 by LR Clonase II. These constructs were transformed into WT *Arabidopsis* or GNL1-Myc BFA-sensitive *Arabidopsis* line^21^ by floral dipping using *Agrobacterium tumefaciens*^54^. Double fluorescent lines were obtained by crossing of the lines above. For all observations, seeds were treated with 0.1% Triton in water during 5 minutes, then washed three times with sterile deionized water and kept 3-4 days at 4°C in water. The seed were then sterilized using a 0.9% chlorine/0.1% Triton solution during 20 minutes and sown on ½ Murashige and Skoogs (MS) agar medium plates (0.8% plant agar, 1% sucrose, and 2.5mM morpholinoethanesulfonic acid pH5.8 with KOH). The seedlings were grown 5 days vertically in long day conditions (150 mE/m^2^/s -1, 16h light/ 8h dark) at 22°C.

### Live cells airyscan and confocal microscopy

Live time-lapse and Z-stacks acquisitions in Fig. 1, 2, 3 (except for 3i-n), 5, 6 and extended data 1e-h, 2, 3a-g, 5, 6 were done using the Airyscan1 module of a Zeiss LSM 880 using 40X oil-immersion objective (NA 1.3). Live cell airyscan microscopy were performed on time-lapse acquisition for the dynamic tracking of compartments association, 3D-reconstruction using the IMARIS software and cargo trafficking analysis using the Retention Using Selective Hooks (RUSH) system. In all cases, seedlings were mounted between an Epredia™TM SuperFrost Plus slide and a 20×20 mm #1.5 coverslip spaced by double-sided tape (except for RUSH observation, the double-sidedtape was not used), in this small chamber the seedlings were incubated with ½MS liquid medium (1% sucrose, and 2.5mM morpholinoethanesulfonic acid pH5.8 with KOH). Dynamic time-series acquisitions were performed as follow: 150 frames, 1 frame every 2 seconds. 3D Z-stacks series were done with a Z-step of 100 nm through 10 µm thickness. The 561 nm excitation laser was used for mCherry or mRFP while the 488 nm laser was used to imaged GFP, the MBS used was 488/561/633 and the filters used were BP495-550 + LP570. Live cell confocal imaging in Fig. 3i-n, Extended data 1a-d, 3h-j was performed with a Zeiss LSM 880 using 40X oil-immersion objective (NA 1.3). The 488 nm excitation laser was used for GFP, the 561 nm laser was used for mRFP and the 405 nm laser was used for the tagBFP2.

### 3D Modelling

3D reconstruction and analysis was performed using the IMARIS10 software. Prior to IMARIS analysis a median filter (3×3×1) and a subtract background (rolling radius 2, except for p35S::NAG1-EGFP where no subtract background was done) were applied to the Z-stack acquisitions. 3D volumes were created for both channels and then sorted according to the distance between either mCherry-MEMB12 or mRFP-SYP31 compartments with the nearest p35S::NAG1-EGFP compartment. We then classified them as Golgi-associated or Golgi-independent and calculated their areas.

### Quantitative live tracking of compartment association

The dynamic analysis was done using the Detect, Track, and Colocalize (DTC) plugin of ImageJ (https://github.com/fabricecordelieres/IJ-Plugin_DTC)^27^. Images were pre-treated as follow before being analysed with the DTC: individual cells were manually delimited using the polygon selection tool and added to the ROI manager; a Gaussian blur filter (sigma 2) and a subtract background (rolling radius 10, except for p35S::NAG1-EGFP were the rolling radius was 5) were applied to all images. The geometrical centres or centroids were calculated for all compartments in the two channels and for each time points. The compartments were tracked if they were at least detected in two successive frames. The maximum of displacement of a given object between two time-frames was setup to 9 pixels. We setup a reference value for the distance of association between two compartments by measuring, for each couple of markers, the average distance between two compartments when they were clearly associated. This distance of association was then applied to the time-lapse acquisitions to distinguish when a compartment in the first channel is associated to another in the second channel. If the distance between one compartment in the first channel and the nearest compartment in the second channel is inferior to the reference distance of association, then we considered that they were associated. Oppositely, if their distance was superior to the reference distance of association, then we considered that they were not associated, e.g. independent. For the RUSH quantification, we applied as association distance the resolution limit for the AiryScan cell per cell analysed. All values were calculated for each detected compartment, in both channels and each time-frames, the values were then sorted using an excel macro we established and made available on demand.

### Inducible cargo release using the Retention Using Selective Hooks (RUSH) system

Cloning of the RUSH system for plant expression was based on multisite Gateway® three-fragment vector construction. In the first vector (pDONR P4-P1r) we introduced the mild-expressing UBQ10 promoter (pUBQ10)^24^, in the second cassette (pDONR P1-P2) we introduced the cargo and in the third cassette (pDONR P2r-P3) we introduced the ER-hook. The three cassette were combined into the pB7m34GW plant expression vector. The luminal cargo sequence was made of an ER-signal peptide (SP) followed by EGFP (optimized for Arabidopsis) and FKBP (Table 1). The ER-hook sequence was made of a peptide 2A (P2A), an ER-signal peptide (SP) followed by TagBFP2, a linker, FKBP and the ER-retention HDEL peptide (Table 1). The cargo and hook sequences were obtained by gene synthesis and then integrated into pDONR P1-P2 and pDONR P2r-P3, respectively. We transformed by floral dip the pUBQ10::SP-EGFP-FKBP-P2A-SP-TagBFP2-linker-FKBP-HDEL/ pB7m34GW construct into Arabidopsis thaliana plants expressing the pUBQ10::mCherry-MEMB12 marker. We selected 3 homozygote mono-insertion lines and further worked with one line. The selection was based on resistance to Basta (10 µM phosphinotricine final concentration in MS plate). Live imaging microscopy was performed on 5 days old seedlings that were mounted between a slide and coverslip into ½ MS liquid media containing 200 µM of the synthetic FKBP ligand GPI1485 (MedChemExpress). The glass slides were then vacuumed 90 seconds at 600mmHg and then immediately placed under the microscope for acquisition. The co-localization quantification was based on the limit of resolution measured in our airyscan images, the geometrical centres or centroids were calculated for all compartments in the two channels. If the distance between one compartment in the first channel and the nearest compartment in the second channel is inferior to 166 nm, then we considered that they were co-localized. Oppositely, if their distance was superior to 166 nm, then we considered that they were not co-localized.

### Inhibitor treatments

Metazachlor (Mz, Merck, Sigma PESTANAL®) treatment was performed on seedlings grown for 5 days on 100 µM Mz-containing ½ MS plates. Fumonisin B1 (FB1, Merck Sigma) treatment was performed on 5-day old seedlings grown on normal ½ MS plates that were transferred for 16 hours into multiwall plates containing liquid ½ MS medium with 2.5 µM FB1. Brefeldin A (BFA, Merck Sigma) treatment was performed on seedlings grown for 5 days on ½ MS plates that were transferred for 1.5 hours into multiwell plates containing liquid ½ MS medium with 50 µM BFA.

### Immuno-staining

The immuno-localization protocol was adapted from Boutté and Grebe, 2014^55^ and used an Intavis InsituPro VSi robot. After seedling fixation (4% paraformaldehyde dissolved in MTSB: 50mM PIPES,5mM EGTA, 5mM MgSO4 pH 7 with KOH) and root tip dissection on polylysine-coated Epredia® SuperFrost Ultra Plus Gold slides (ThermoFisher Scientific), the slides with the root samples were mounted under a counter slide using 1X MTSB solution and positioned in the InsituPro VSi holder. The program for single and double staining are described in the supplemental Table 2. At the end of the program, slides were mounted under a coverslip #1.5 using ProlongGold anti-fade mounting-medium (ThermoFischer Scientific). The following antibodies and nanobodies were used: anti-MEMB12^40^ 1/100, goat anti-rabbit-AlexaFluor AF594 1/300 (Abcam), GFPbooster-ATTO647N 1/200 (Proteintech) and RFPbooster-ATTO594 1/200 (proteintech).

### Lifetime tau-Stimulated Emission Depletion (τ-STED) microscopy

Acquisitions were done using a Leica SP8 (DMI6000 TCS SP8 X) equipped with a STED and a FALCON module in sequential mode with a 93X Glycerol immersion objective (NA 1.4). The excitation laser wave lines were 594 nm for the AF594 and ATTO594 and 640 nm for the ATTO647N. The power of the excitation laser was set between 5-15%, the power of the 775 nm depletion laser (used for AF594, ATTO594 or ATTO647N) was set between 30-50%, depending on the expression strength of the protein marker and the fluorophore used. Emission signals were collected between 610-640 nm for AF594/ATTO594 and between 655-690 nm for ATTO647N with a frame accumulation between 10 and 12 for FLIM measurements. Images were automatically processed by the τ-STED module of Leica.

### Morphometric analysis of τ-STED images

τ-STED images were filtered using a median filter (3×3×2) and converted into binary images. Then the analyse particle plugin of ImageJ was applied using the following set up: area from 0.04 to infinity, circularity from 0 to 1, add to ROI manager and exclude on edge. The ROIs were applied on the original images and ROIs that do not correspond to an individual structure were eliminated. Measurement were set up to obtain the area and shape descriptor of each individual ROI, from the shape descriptor we extracted the circularity; a line has a circularity of 0 and a circle a circularity of 1; and the solidity. The solidity is a ratio of the full convex area on the area of an object, if the full convex area and the area are the same the solidity will be close to 1.

### Expansion Microscopy (ExM)

The proExM method was adapted from Tillberg et al., 2016. After fixation and immunolabeling root tips are glued to 12 mm round glass coverslips coated with Poly-L-Lysine prior to proExM protocol. Then after digestion, gels containing the roots are detached from the coverslip and carefully dissected to eliminate excess of empty gel. Root-containing gels are transferred to a clean container and expanded in water. Water is renewed several times until plateau is reached. Gels are finally mounted in a coverslip glass previously treated with poly-L-lysine for imaging. Expended root tips were imaged using a Leica SP8 (DMI6000 TCS SP8 X) in sequential mode with a 25X water plunging objective (NA 0.8). The excitation laser wave lines were 594 nm for the AF594 and 647 nm for the ATTO647N. The power of the excitation laser was set between 5-15%, and emission signals were collected between 610-640 nm for AF594 and between 655-690 nm for ATTO647N.

### Calculation of the limit of resolution

References were taken in images from τ-STED, ExM and airyscan images to calculate the resolution of the system directly on the image using FIJI. The intensity distributions were fitted to the nearest gaussian and the full width at half maximum “d” value were used to calculate the resolution with the following formula:

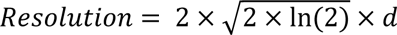

A minimum of 20 measures have been done for each technics and “*d*” values were used only if the fitting curve displayed a R^2^>0,90.

### Statistical information

The normality test was performed on each data set by the Shapiro-Wilk test to determine whether the data follow a Gaussian distribution. To compare two groups, unpaired t-test’s rank sum test was used if the datasets follow a Gaussian distribution (with welch’s correction if the standard deviation of the two datasets are separated by more than one unit). If the datasets do not follow a Gaussian distribution, the two-sided Wilcoxon’s rank sum test was used. To compare three groups or more, the One-Way ANOVA Brown-Forsythe test was performed if all the datasets follow a Gaussian distribution, otherwise the Kruskal-Wallis test was used.

---

**Extended data 1.**
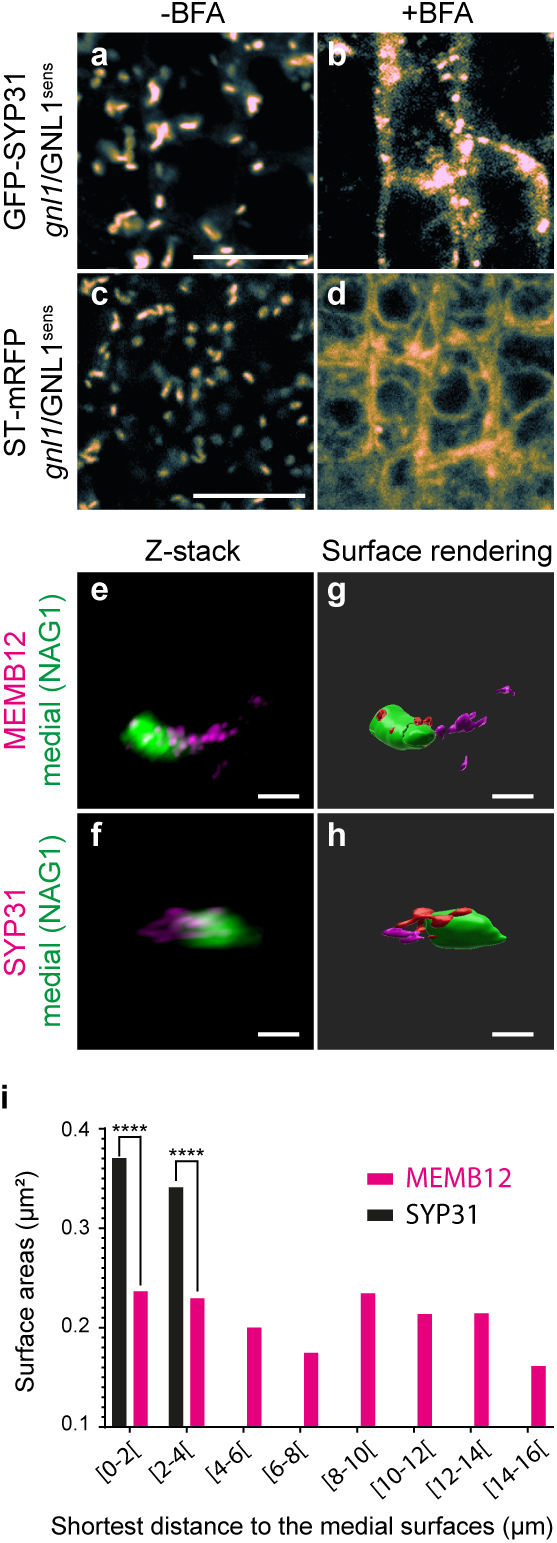
The SYP31-positive Golgi Entry Core Compartment (GECCO) exists in Arabidopsis root epidermal cells and is mostly associated to the Golgi in untreated cells. (**a-d**) Arabidopsis epidermal root cells expressing either GFP-SYP31 (**a, b**) or ST-mRFP (**c, d**) in the gnl1 mutant background complemented with a BFA-sensitive version of GNL1. (**a, c**) In absence of BFA, SYP31 and ST localize to Golgi-like structures. (**b, d**) In presence of BFA, SYP31 (**b**) remains in dotty-structures while ST (**d**) is redistributed to the ER-network. (**e-h**) Additional display of Fig. 1g-n. 3D acquisition (**e, f**), reconstruction and IMARIS surface modelling (**g, h**) of either mCherry-MEMB12 x NAG1-EGFP (**e, g**) or mRFP-SYP31 x NAG1-EGFP (**f, h**). MEMB12 is more fragmented and distant from the medial-Golgi than SYP31. (**i**) Another way of representing the data of Fig. 1p. Quantification of the areas generated by surface rendering. SYP31-structures are close to the Golgi (within 4 µm) and bigger than MEMB12-structures that are distributed at various distances from the Golgi. A subpopulation of MEMB12 lies within 4µm-distance from the Golgi while another subpopulation is located between 8-14 µm-distance from the Golgi (n=27 015 or n=3 727 compartments for MEMB12 x NAG1 or SYP31 x NAG1, respectively, out of 12 cells for each markers. **** p<0.0001 by two-sided Wilcoxson rank sum test.). Scale bars are 10 µm in **a-d** and 1µm in **e-h**.

**Extended data 2.**
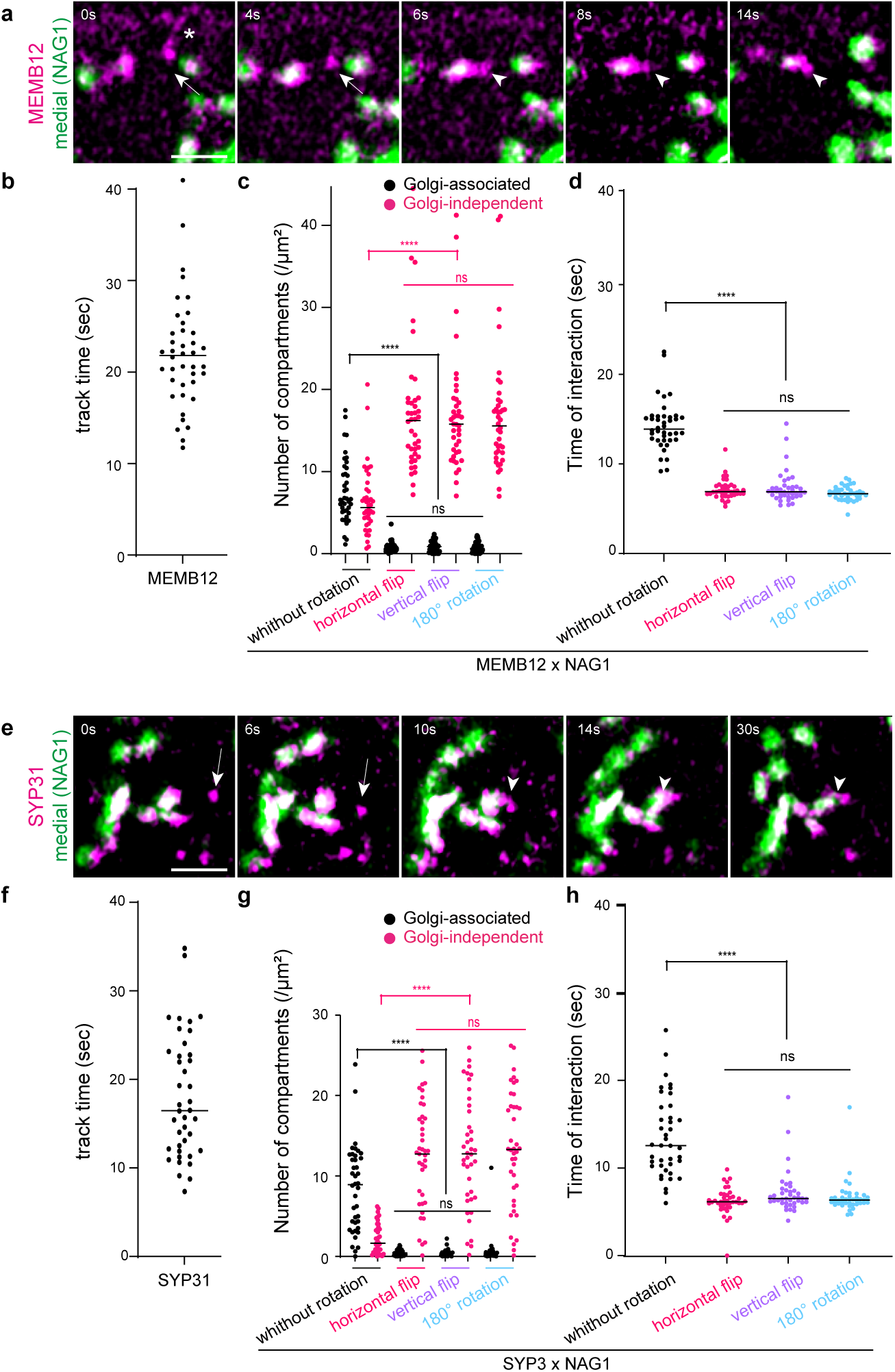
Validation of the quantitative data obtained from the time-lapse airyscan acquisition of Fig. 2. (**a, e**) Time-lapse airyscan acquisition in root epidermal cells of either (**a**) mCherry-MEMB12 x NAG1-EGFP (**a**) or mRFP-SYP31 x NAG1-EGFP (**e**). The time series show an independent MEMB12- or SYP31-compartment that undergo an association with the medial-Golgi. The white star in a indicates a tubular structure. (**b, f**) Upper edge control: individual MEMB12- (**b**) or SYP31- (**f**) compartments could be tracked for an average track time of 22 sec and 17 sec, respectively. (**c, g**) Lower edge control: one channel was flipped either horizontally or vertically, or rotated by 180°. Without rotation or flip, the number of MEMB12-compartments (**c**) that either remain associated to or remain independent from the medial-Golgi is similar, contrastingly to SYP31 (**g**) that mostly associate with the Golgi. With rotation or flip of one channel, the number of MEMB12- or SYP31-compartments that remain associated to the Golgi drop to close to 0 while the number of compartments that remain independent from the Golgi drastically increases. (**d, h**) Time of interaction between either MEMB12 (**d**) or SYP31 (**h**) with the medial-Golgi. Without rotation or flip, the average interaction time is around 15 sec for MEMB12/medial-Golgi and 12 sec for SYP31/medial-Golgi, respectively. With rotation or flip, the average time of interaction drops to 5-6 sec for both MEMB12 and SYP31. n=40 cells for all set of data, **** p<0.0001, ns p<0.1234 by kruskal-wallis and two-sided Wilcoxon’s rank sum tests. All scale bars are 1µm.

**Extended data 3.**
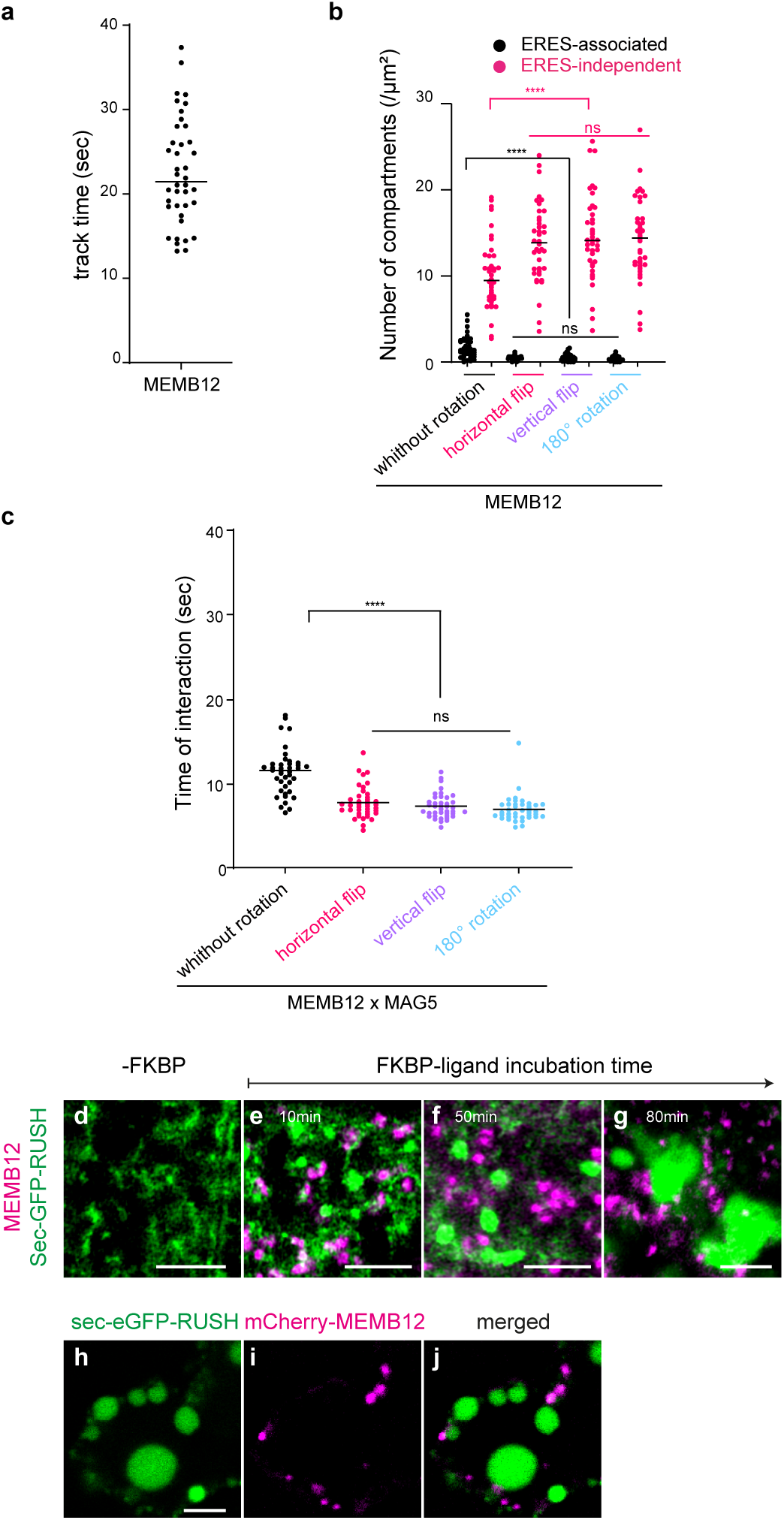
Validation of the quantitative data obtained from the time-lapse airyscan acquisition of Fig. 3 and additional RUSH-images. (**a**) Upper edge control: individual MEMB12-compartments could be tracked for an average track time of 22 sec. (**b**) Lower edge control: one channel was flipped either horizontally or vertically, or rotated by 180°. Without rotation or flip, the number of ERES-independent MEMB12-compartments is much higher than the number of ERES-associated compartments. With rotation or flip of one channel, the number of MEMB12-compartments that remain associated to the ERES drop to close to 0 while the number of compartments that remain independent from the ERES drastically increases (n=40 cells, **** p<0.0001, ns p<0.1234 by either one-way ANOVA Broth-Forsythe and two-sided t-test or Wilcoxon’s rank sum test for ERES-independent or ERES-associated MEMB12, respectively). (**c**) Time of interaction between MEMB12-compartments and the ERES. Without rotation or flip, the average interaction time is around 12 sec. With rotation or flip, the average time of interaction drops to 5-6 sec (n=40 cells, **** p<0.0001, ns p<0.1234 by kruskal -wallis and two-sided Wilcoxon’s rank sum tests). (**d-g**) Confocal images of root epidermal cells stably expressing both sec-GFP-RUSH construct and mCherry-MEMB12. (**d**) In absence of FKBP ligand, sec-GFP-RUSH localizes to the ER. In presence of FKBP-ligand, sec-GFP localizes to small dotty structures as well as bigger pro-vacuole-like compartments after 10 min of incubation, or bigger vacuoles after 80 min of incubation. (**h-j**) Transient expression of the sec-GFP-RUSH (**h**) construct in epidermal cotyledon cells stably expressing mCherry-MEMB12 (**i**, merged in **j**). Vacuole-like compartments are labelled by sec-GFP-RUSH upon incubation with FKBP-ligand. All scale bars are 4µm.

**Extended data 4.**
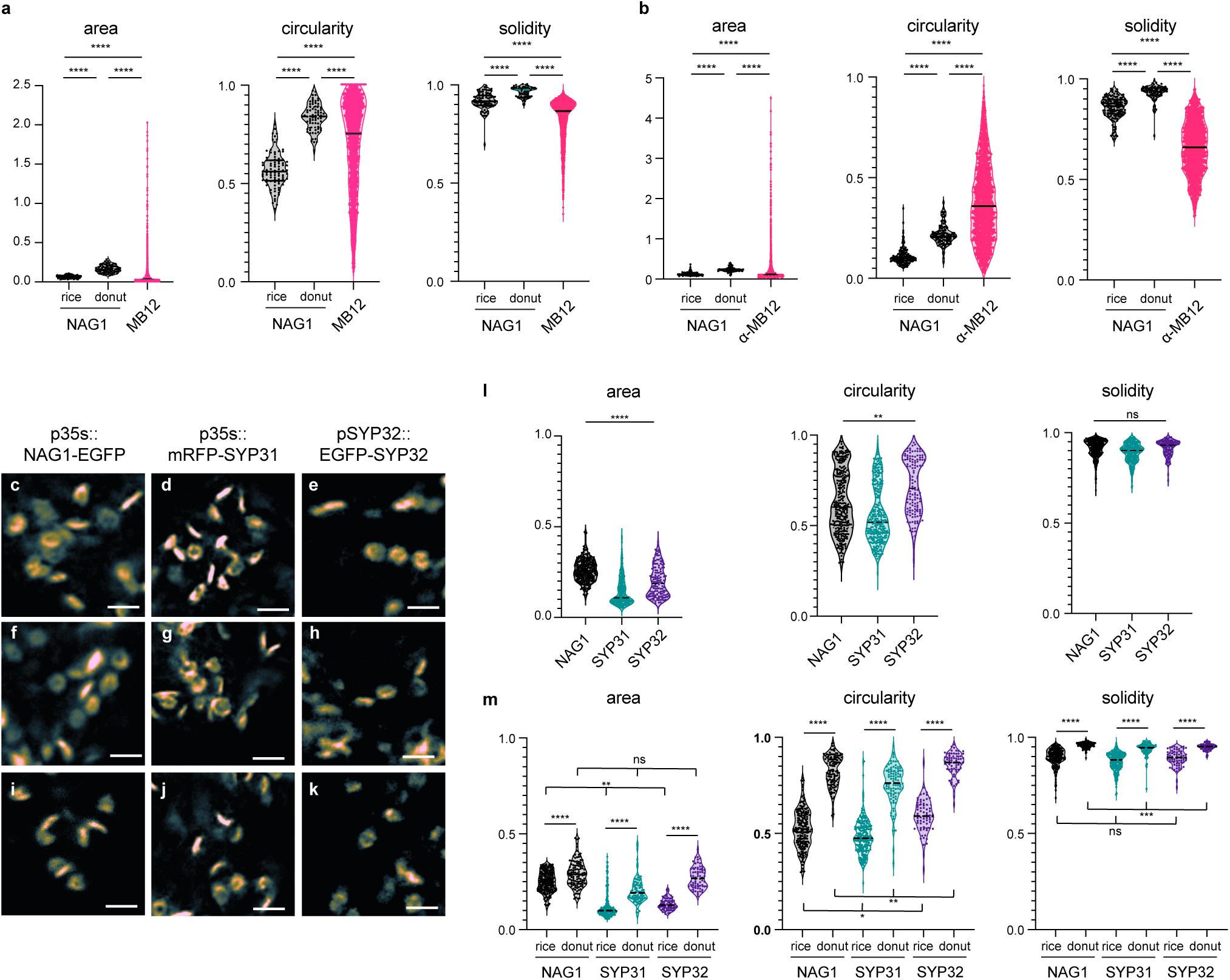
MEMB12 labels a heterogeneous network while NAG1, SYP31 and SYP32 label cisternae. (**a, b**) Morphometric analysis of either mCherry-MEMB12 x NAG1-EGFP (**a**) or α-MEMB12 immunostaining in NAG1-EGFP expressing plants (**b**). MEMB12-structures display higher area and lower circularity and solidity with a more spread repartition of the values than NAG1. The binomial distribution of NAG1 values in Fig. 4 is explainable by the orientation of the cisterna during the acquisition. If the cisterna is oriented on the side it will look like a rice grain. If the cisterna is viewed from the top or bottom it will look like a donut. Donut-like structures have an area, circularity and solidity index higher than rice grain structures. (**c-k**) τ-STED images of either NAG1-EGFP (**c, f, i**), mRFP-SYP31 (**d, g, j**) or EGFP-SYP32 (**e, h, k**). (**l, m**) Morphometric analyses of images in **c-k**. Although being all localized on cisternae-like structures, SYP31- and SYP32-compartments are different from the medial-Golgi labelled by NAG1. Notably, the area of SYP31- and SYP32-compartments is smaller than for the NAG1 medial-Golgi. n= 315 compartments including 199 “rice” and 116 “donut” for NAG1, n=311 compartments including 217 “rice” and “90” donuts for SYP31, n= 134 compartments including 68 “rice” and 66 “donut” for pSYP32 from 16 cells. * p<0.05; ** p<0.005; *** p<0.0005; **** p<0.0001 by Wilcoxon rank sum test for “rice” and “donut” comparisons (**a, b, m**) and Brown Forsythe Anova (**a, b, i, m**). Scale bars are 1µm.

**Extended data 5.**
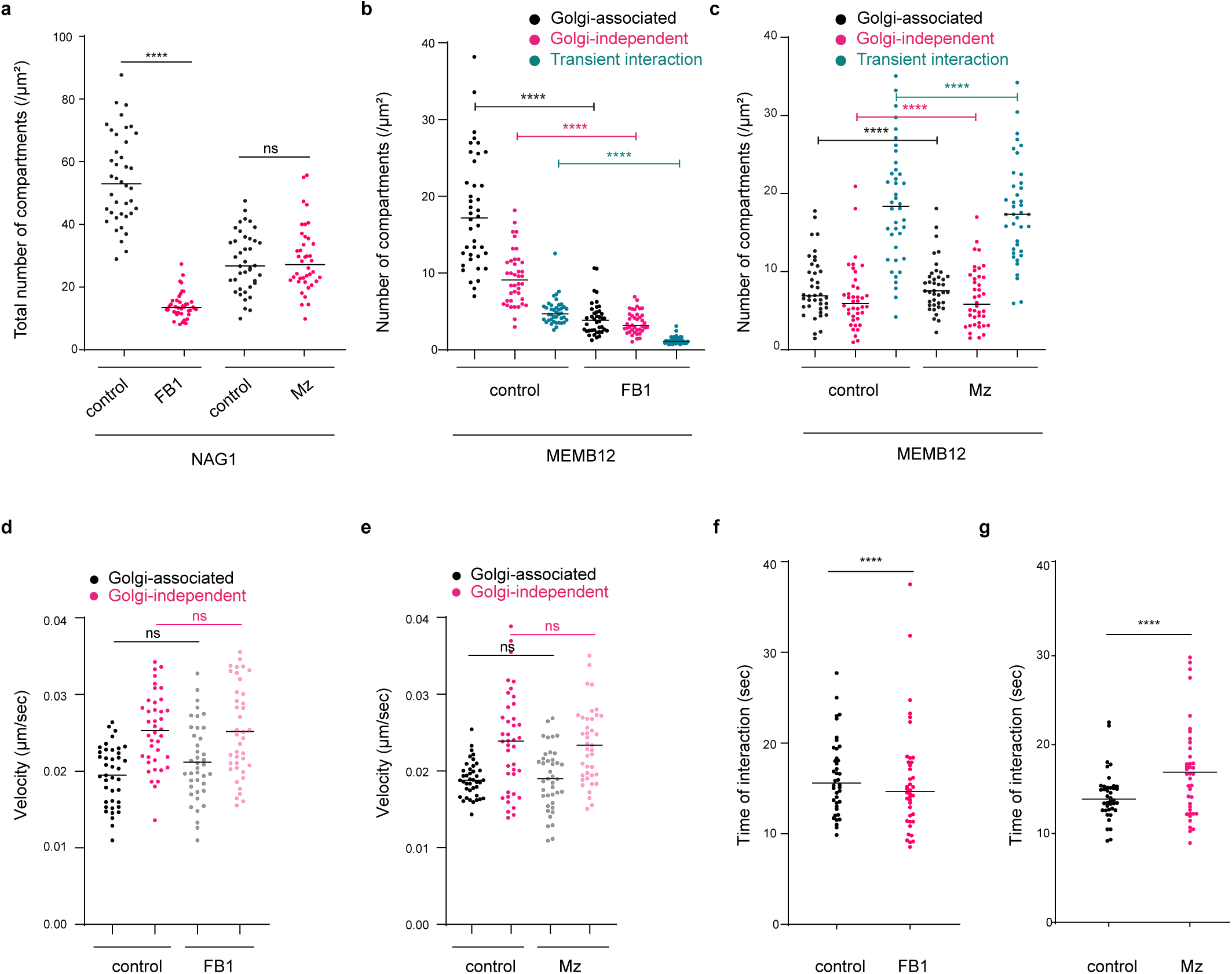
Sphingolipid function in the MEMB12-compartments dynamic interaction with the medial-Golgi (additional data and controls from Fig. 5). (**a**) Quantification of the total number of NAG1 medial-Golgi showing a severe decrease upon FB1 but not upon Mz treatment (n=40 cells, **** p<0.0001 by either two-sided Wilcoxon or t-test rank sum test for FBI or Mz treatment, respectively). (**b, c**) Number of Golgi-associated, Golgi-independent or transient interaction of MEMB12 with the medial-Golgi upon either FB1 (**b**) or Mz (**c**) revealing a major drop of all compartments upon FB1 but not Mz (n=40 cells, **** p<0.0001 by either two-sided Wilcoxon or t-test (welch correction) rank sum test for FBI or Mz treatment, respectively). (**d, e**) The velocity of the MEMB12-compartments that are either associated with or independent from the medial-Golgi is not altered in either case nor by FB1 nor by Mz treatment (n=40 cells, ns p<0.1234 by two-sided t-test (welch correction) rank sum test). (**f, g**) The time of interaction between MEMB12-compartments and the medial-Golgi is not altered upon FB1 and is slightly increased upon Mz (n= 40 cells, **** p<0.0001 by two-sided t-test (welch correction for Mz) rank sum test).

**Extended data 6.**
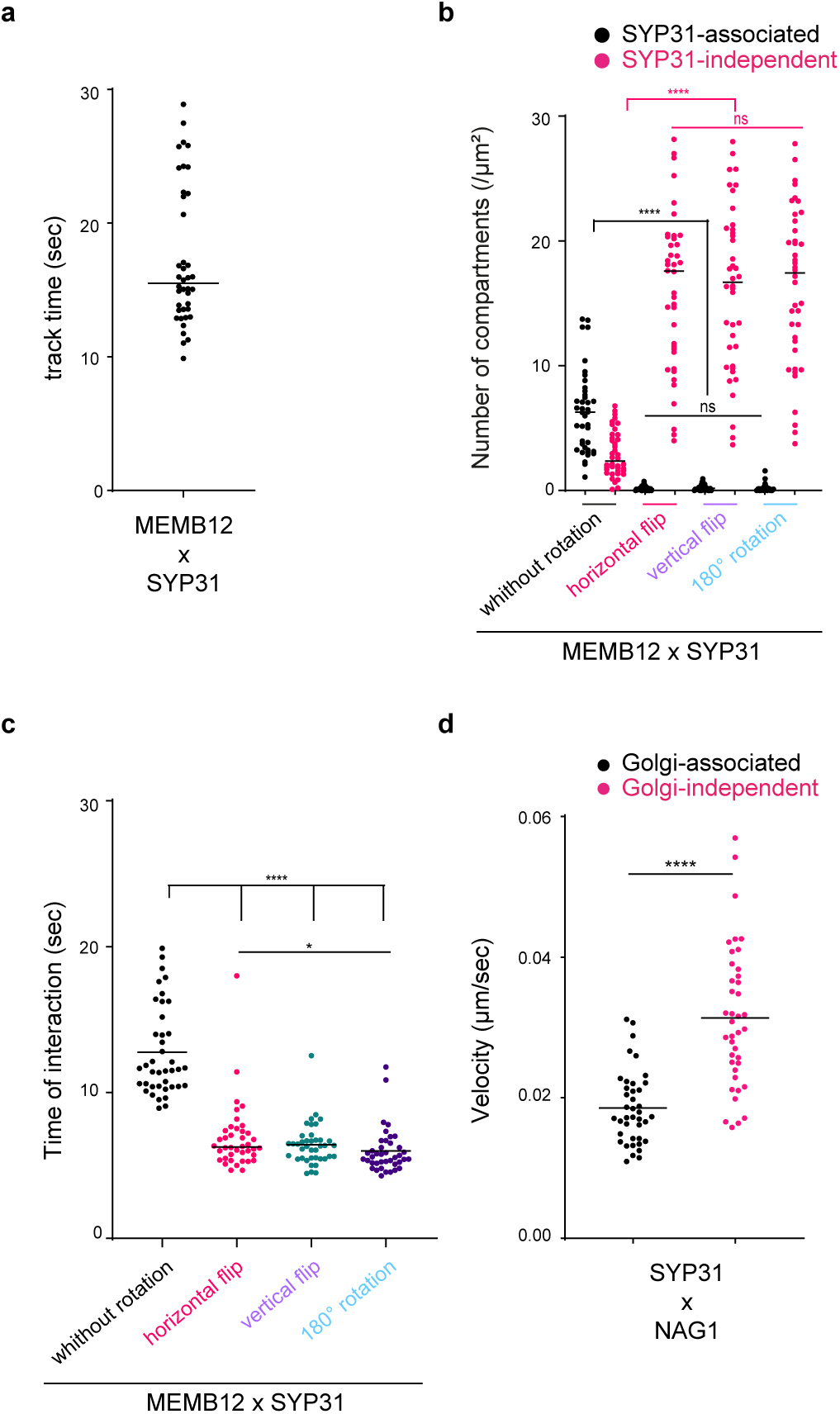
Validation of the quantitative data obtained from the time-lapse airyscan acquisition of Fig. 6. (**a**) Upper edge control: individual MEMB12-compartments could be tracked for an average track time of 15-17 sec (n=40 cells, **** p<0.0001 by two-sided Wilcoxon or t-test rank sum tests respectively for FBI and Mz treatment. (**b**) Lower edge control: one channel was flipped either horizontally or vertically, or rotated by 180°. Without rotation or flip, the number of SYP31-independent MEMB12-compartments is much higher than the number of SYP31-associated compartments. With rotation or flip of one channel, the number of MEMB12-compartments that remain associated to SYP31-compartments drop to close to 0 while the number of compartments that remain independent from SYP31-compartments drastically increases (n=40 cells, **** p<0.0001, ns p<0.1234, by kruskal-wallis and Wilcoxon’s rank sum tests). (**c**) Time of interaction between MEMB12- and SYP31-compartments. Without rotation or flip, the average interaction time is around 12 sec. With rotation or flip, the average time of interaction drops to 5-6 sec (n=40 cells, **** p<0.0001, * p<0.0332 by kruskal-wallis and Wilcoxon’s rank sum tests). (**d**) The velocity of SYP31-compartments is decreased when it associates to the medial-Golgi. n=40 cells, **** p<0.0001, by Wilcoxon’s rank sum test.

